# Geissoschizine scaffolding enzymes shape monoterpene indole alkaloid biosynthesis

**DOI:** 10.64898/2026.04.29.721795

**Authors:** David Brelu-Brelu, Enzo Lezin, Emma Parker Miller, Duchesse-Lacours Zamar, Mickael Durand, Sitakanta Pattanaik, Barunava Patra, Nicolas Gautron, Caroline Birer Williams, Thomas Perrot, Arnaud Lanoue, Audrey Oudin, Pierre Le Pogam, Mehdi A Beniddir, Nathalie Giglioli-Guivarc’h, Benoit St-Pierre, Georges Massiot, Nicolas Papon, Ling Yuan, Sijin Li, Chao Sun, Sébastien Besseau, Vincent Courdavault

## Abstract

For decades, medicinal plants have been an invaluable source of therapeutics for human health. Among plant specialized metabolites, monoterpene indole alkaloids (MIAs) display remarkable chemical diversity and potent bioactivities, including anticancer properties. While the biosynthetic pathways leading to major MIAs such as vincristine, vinblastine, and camptothecin have been extensively studied, the mechanisms regulating metabolic flux within these pathways remain poorly understood. Here, we uncover an unexpected regulatory layer in MIA biosynthesis involving medium-chain dehydrogenase/reductase (MDR) proteins. Using a combination of *in vitro* biochemical assays, pathway reconstitution in planta, protein–protein interaction analyses, and metabolic engineering in yeast, we show that specific MDRs do not function as classical catalysts but instead enhance both the activity and diversify the stereochemical outputs of geissoschizine synthase (GS). These proteins physically interact with GS and strictosidine β-glucosidase (SGD), forming ternary complexes that likely facilitate substrate channeling of reactive intermediates. Importantly, introduction of these MDRs into engineered yeast strains leads to a dramatic increase in the production of geissoschizine, a central MIA precursor. Collectively, our findings reveal a previously unrecognized role for MDR proteins as regulators of metabolic flux and highlight their potential as powerful tools for metabolic engineering of valuable plant natural products.

## INTRODUCTION

Monoterpene indole alkaloids (MIAs) are a complex class of natural products mainly synthesized by plants of the Apocynaceae and Rubiaceae families. With more than 3,000 compounds identified to date, MIAs display an extraordinary structural diversity that underlies their wide range of biological activities and roles *in planta*. Although MIAs primarily contribute to plant adaptation to the environment, notably through defense against biotic aggressors, they are also extensively employed in human medicine, as exemplified by the anticancer drugs vinblastine and vincristine from the Madagascar periwinkle (*Catharanthus roseus*) or camptothecin from the happy tree (*Camptotheca acuminata*). Remarkably, this extensive chemical diversity is associated with highly variable biosynthetic outputs across species, yet it arises from a surprisingly limited set of conserved enzymatic steps.

All MIAs indeed derive from the common precursor strictosidine, the first committed MIA, which results from the Pictet–Spengler condensation of tryptamine and secologanin catalyzed by strictosidine synthase (STR) (**Fig. 1**). Strictosidine is subsequently deglucosylated by strictosidine β-glucosidase (SGD), yielding a highly reactive aglycone that spontaneously rearranges into 4,21-dehydrogeissoschizine. This unstable intermediate undergoes non-enzymatic cyclization to form cathenamine and 19-*epi*-cathenamine, as previously reported ^1,2^. This dynamic ensemble of hemiaminal, enamine and iminium species constitutes a central branching point for downstream transformations catalyzed by medium-chain dehydrogenases/reductases (MDRs) ^3,4^. MDRs involved in MIA metabolism selectively reduce iminium intermediates derived from the strictosidine aglycone, thereby channeling metabolic flux towards distinct alkaloid scaffolds. For instance, tetrahydroalstonine synthases (THAS) catalyze the stereoselective 1,2-iminium reduction of cathenamine, leading to the formation of the heteroyohimbane tetrahydroalstonine (THA) ^5,6^. In a closely related reaction, heteroyohimbane synthase (HYS) reduces both cathenamine and 19-*epi*-cathenamine, yielding a mixture of heteroyohimbane MIAs, including tetrahydroalstonine, ajmalicine and mayumbine^6^. A similar, yet strategically distinct, function is fulfilled by geissoschizine synthase (GS), which catalyzes the reduction of the conjugated iminium of 4,21-dehydrogeissoschizine to form the secoyohimbane 19*E*-geissoschizine^7^. Such a reaction occupies a central position in MIA biosynthesis, as 19*E*-geissoschizine constitutes a common precursor for several major alkaloid scaffolds, including aspidosperma (vindoline), iboga (catharanthine), and sarpagan (ajmaline), depending on the plant species. In addition, yohimbane synthase (YOS) performs the cyclization of the 4,21-dehydrogeissoschizine iminium and subsequently reduces the resulting intermediate into four yohimbane stereoisomers including rauwolscine, yohimbine, and corynanthine ^8^. Interestingly, this yohimbane synthesis can also proceed through a distinct route involving a first GS-catalyzed reduction of 4,21-dehydrogeissoschizine combined with a reduction/cyclization catalyzed by a second MDRs, including MSTRG.5530, MSTRG.5531 and MSTRG.5534, in *Rauvolfia tetraphylla* ^8^.

**Fig. 1.**
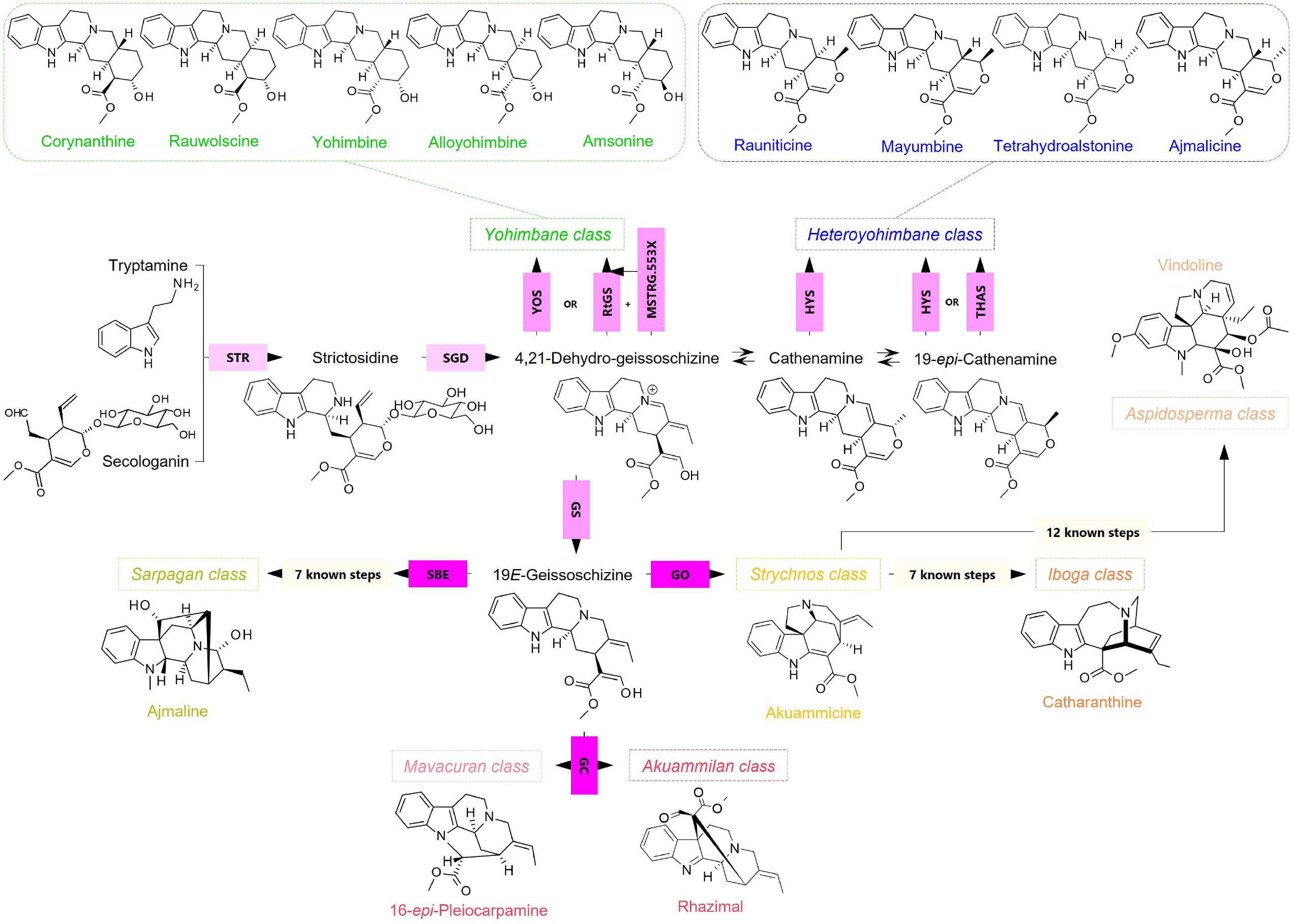
Biosynthetic pathways of monoterpene indole alkaloid (MIA) classes. Pink boxes denote each enzyme characterized to be involved in MIAs biosynthesis. The pink gradient indicates functional categorization: pale pink corresponds to enzymes responsible for the formation of the common and central MIA precursor strictosidine, whereas medium and dark pink highlight enzymes driving the diversification of MIA scaffolds. The enzymes shown in medium pink belong to the medium-chain dehydrogenases/reductases protein family and proteins outlined in dark pink are members of the cytochrome P450 family. STR: strictosidine reductase. SGD: strictosidine β-*D*-glucosidase. YOS: Yohimbane synthase. GS: Geissoschizine synthase. MSTRG.553X encompasses three orthologous proteins (MSTRG.5530, MSTRG.5531 and MSTRG.5534) previously identified in *R. tetraphylla*. HYS: Heteroyohimbane synthase. THAS: Tetrahydroalstonine synthase. GS: Geissoschizine synthase. SBE: Sarpagan bridge enzyme. GO: Geissoschizine oxidase. GC: Geissoschizine cyclase.

Throughout the MIA biosynthetic pathway, these MDR products are subsequently metabolized by several enzymes from distinct families to generate the whole diversity of MIAs accumulated *in planta*. This usually starts with reactions catalyzed by cytochromes P450 (P450s) from the CYP71 clan including distinct oxidative cyclizations of 19*E*-geissoschizine by geissoschizine oxidase (GO) initiating the synthesis of strychnos/aspidosperma MIAs ^7^, geissoschizine cyclase (GC) producing strychnos, akuammilane and mavacurane MIAs ^9^or sarpagan bridge enzyme (SBE) for sarpagan MIA synthesis ^10^. Besides cyclisation, CYP71 also catalyzes aromatization of geissoschizine but also heteroyohimbane and yohimbane as reported for alstonine synthase (AS), serpentine synthase (SS) and the whole set of MIA-associated CYP71 ^11–13^. Several other dehydrogenases/reductases catalyze downstream reactions along the pathway including redox 1, redox 2 or dihydroprecondylocarpine acetate synthase (DPAS) for the synthesis of stemmadenine ^14,15^or vomilenine reductase 2 (VR2) for ajmaline ^16,17^ besides the other reductases from the pre-strictosidine branch, notably including the 10-hydroxygeraniol oxidoreductase (10HGO) ^18^. Lastly, P450s, methyltransferases and dioxygenases contribute to the radiating diversification of MIAs ^19^.

While many enzymes of the MIA pathway have been characterized across different plant species, the mechanisms that shape MIA biosynthesis and diversification remain largely unresolved. Increasing evidence nevertheless points to protein–protein interactions (PPIs) as an important process of biosynthesis regulation. In *C. roseus* and *Mitragyna speciosa*, both THAS and HYS interact with SGD in the nucleus, thus forming a metabolon that promotes the efficient conversion of strictosidine aglycone into cathenamine and channels metabolic flux towards heteroyohimbane MIAs ^5,6,20^. In contrast, to date, no comparable interaction between GS and SGD has been reported to trigger geissoschizine biosynthesis, even though geissoschizine-derived MIAs represent the most abundant alkaloids *in planta*, as exemplified by catharanthine and vindoline in *C. roseus*. Moreover, whereas multiple THAS-encoding genes are typically present in MIA-producing plant genomes, GS is often represented by only one ^21,22^ or several gene copies ^23– 25^. This suggests that additional mechanisms may contribute to the control of metabolic flux through the geissoschizine branch.

To get insights into this potential process, we leveraged the multi-omics resources recently generated for the medicinal plant *Tabernaemontana elegans* to investigate the functional diversity of MDRs ^22^. Using a combination of *in vitro* biochemical assays, pathway reconstitution *in planta*, analysis of protein–protein interactions, and metabolic engineering in yeast, we show that specific MDRs do not act as classical catalysts but rather enhance both the activity and stereochemical output of GS. We also demonstrate that Geissoschizine Scaffolding Enzymes (GSEs) physically interact with GS and SGD, to form ternary complexes that are likely to promote substrate channeling. Finally, we show that introducing GSEs into engineered yeast strains dramatically improves geissoschizine production, highlighting their potential for metabolic engineering applications.

## RESULTS

### Predicting MDR encoding genes involved in MIA biosynthesis

We used the available toad tree *T. elegans* genome ^22^ to identify MDR-encoding genes involved in MIA biosynthesis through a homology-based screening approach. MDR enzymes related to MIA biosynthesis were used as queries (**Supplementary Table 1**) for BLASTp searches (pident > 60%; qcov > 90%). The 24 putative MIA-associated MDR protein sequences were aligned and analyzed phylogenetically using a neighbor-joining method with 1,000 bootstrap replicates (**Fig. 2**). The resulting tree resolved these candidates into 6 distinct clades corresponding to characterized MDRs. Five proteins (Tel01G.20889, Tel01G.47, Tel01G.56, Tel01G.69, Tel01G.22311) clustered with the THAS-like clade. Among them, four sequences (Tel01G.47, Tel01G.56, Tel01G.69, Tel01G.22311) formed an uncharacterized sub-clade specific to *T. elegans*. One candidate, Tel01G.712, was placed within the GS clade, supporting its putative functional assignment. Another set of four sequences (Tel01G.717, Tel01G.716, Tel01G.2847, Tel01G.24838) grouped with Cr10HGO/RtMSTRG.5534, but Tel01G.715 is highly divergent within the clade. Four sequences (Tel01G.2900, Tel01G.2901, Tel01G.2902, Tel01G.2907) grouped within a CrDPAS-related sub-clade, while four VR2-related genes (Tel01G.2892, Tel01G.2893, Tel01G.2895, Tel01G.2896) formed a separate sub-clade. In both cases, the genes are closely related in sequence (**Supplementary Table 2**) and physically clustered in the genome, consistent with local gene expansion. Finally, three proteins (Tel01G.714, Tel01G.20589, Tel01G.20590) formed the RtMSTRG.5530/RtMSTRG.5531 clade, and the Redox1 group was represented by a single sequence (Tel01G.2903).

**Fig. 2.**
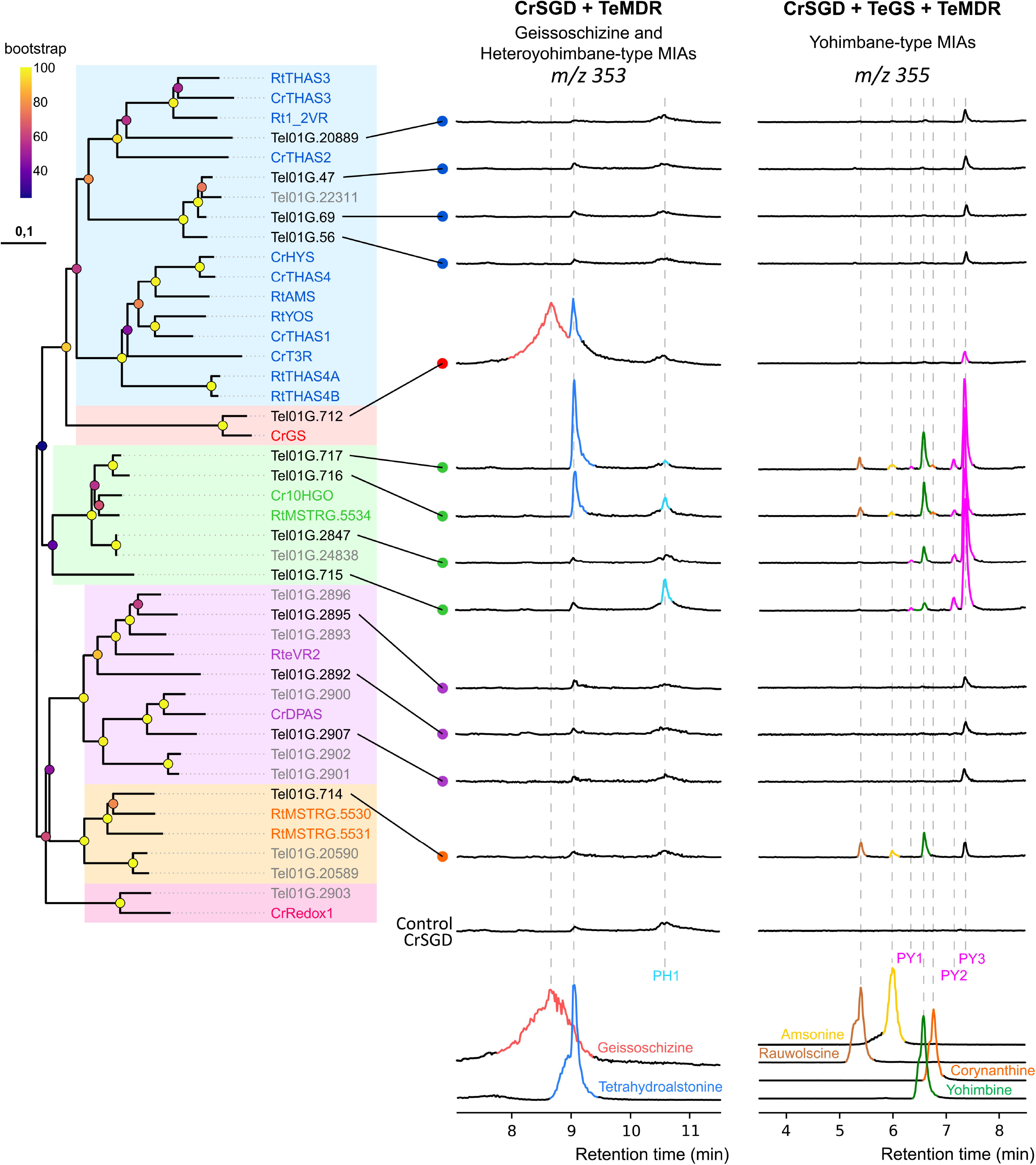
Screening of *T. elegans* MDR candidates by *in vitro* biochemical assays for biosynthesis of geissoschizine, heteroyohimbane-type and yohimbane-type MIAs through direct or GS-associated biosynthetic pathway. Phylogenetic tree illustrating the distribution of MDR-like enzymes from *Tabernaemontana elegans* across clades defined by MIA-related enzymes characterized in *C. roseus* and *R. tetraphylla* (names highlighted in color). TeMDR names shown in black indicate cloned candidates. For enzymatic assays, strictosidine was pre-incubated with purified recombinant SGD from *C. roseus* followed by the addition of MDRs with or without TeGS (left and right panels respectively). Reaction products were analysed by UPLC-MS and compared to authentic standards (through EIC generation at *m/z* 353 and 355). All enzymatic assay chromatograms are shown on the same scale. Colored peaks indicate significant differences compared to the control conditions: CrSGD without TeMDR for geissoschizine and heteroyohimbane production, and CrSGD with TeGS (Tel01G.712) for yohimbane production. PH1, putative heteroyohimbane 1; PY1-3, putative yohimbane 1-3.

### MDRs from T. elegans display heteroyohimbane and yohimbane synthase activities

Among the 24 MDRs identified through homology-based searches, 13 candidates spanning five of the phylogenetic clades were amplified and cloned into the pHREAC expression vector (Tel01G.20889, Tel01G.47, Tel01G.69, Tel01G.56, Tel01G.712, Tel01G.717, Tel01G.716, Tel01G.2847, Tel01G.715, Tel01G.2895, Tel01G.2892, Tel01G.2907, Tel01G.714). Their activities were evaluated following transient expression in *Nicotiana benthamiana* and subsequent protein purification. Recombinant enzymes were incubated with SGD and strictosidine, and reaction products were analysed by UPLC–MS to assess secoyohimbane-, heteroyohimbane-, or yohimbane synthase activities (**Fig. 2**).

Among the tested candidates, only Tel01G.712 exhibited GS activity, consistent with its high sequence identity to GS from *C. roseus* (**Fig. 2**). This enzyme was therefore designated TeGS. Notably, TeGS also catalyzed the simultaneous formation of THA, indicating its capacity to produce both MIA scaffolds. In contrast, none of the candidates homologous to previously characterized THAS enzymes from other plant species displayed detectable heteroyohimbane synthase activity. Of the remaining MDRs, only Tel01G.715, Tel01G.716, and Tel01G.717 catalyzed the formation of THA or a closely related isomer (**Supplementary Fig. 1A-C**), annotated as a putative heteroyohimbane (PH1). These findings suggest that heteroyohimbane synthase activity has emerged independently within distinct yet related MDR clades across MIA-producing plant lineages.

When combined with CrSGD alone, none of the candidates produced yohimbanes except TeGS, which produces traces of the previously reported putative yohimbane 3 (PY3; Stander et al., 2023) (**Fig. 2; Supplementary Fig. 2**). This indicates that no direct orthologue of YOS is present among the 13 selected MDRs. However, as yohimbanes can arise from the concerted action of GS and a second MDR (Stander et al., 2023), each candidate was subsequently assayed in the presence of CrSGD and TeGS. Under these conditions, Tel01G.714, Tel01G.715, Tel01G.716, Tel01G.717, and Tel01G.2847 produced yohimbanes, consistent with their homology to RtMSTRG5530, RtMSTRG5531, and RtMSTRG5534 (**Fig. 2**). Most enzymes predominantly generated PY3, as previously described (Stander et al., 2023), together with additional isomers including rauwolscine, yohimbine, and corynanthine and/or PY1 and PY2. Tel01G.716 and Tel01G.717 exhibited the broadest product profile, producing up to seven distinct yohimbane isomers. These include amsonine, which has the unusual C17R configuration (**Supplementary Fig. 1A-C**). To our knowledge, no other enzyme has been reported to catalyze the formation of this MIA scaffold to date.

Collectively, these results demonstrate that *T. elegans* harbors a set of MDRs exhibiting yohimbane synthase activity when combined with TeGS, in some cases together with a heteroyohimbane synthase activity. Notably, four active MDR genes (Tel01G.714–Tel01G.717) are physically clustered with TeGS (Tel01G.712) in the *T. elegans* genome (**Fig. 3C**). Such genomic organization appears conserved among MIA-producing plants, as supported by microsynteny analyses with *C. roseus* and *R. tetraphylla* (**Supplementary Fig. 3**).

**Fig. 3.**
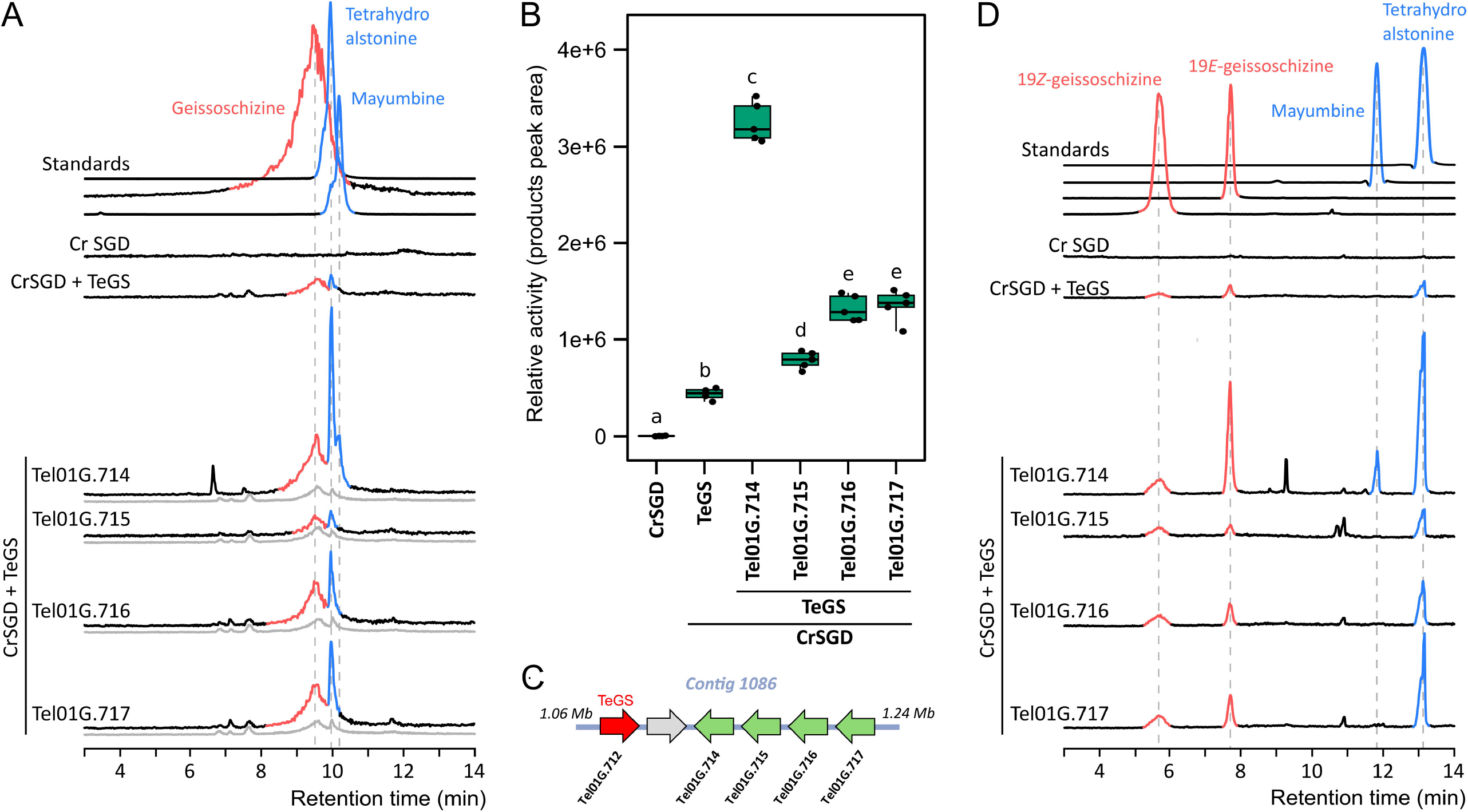
*In vitro* effect of cluster-associated MDRs on TeGS activity in geissoschizine and heteroyohimbane synthesis. Strictosidine was pre-incubated with purified recombinant CrSGD, followed by the addition of TeGS together with Tel01G.714, Tel01G.715, Tel01G.716, or Tel01G.717. CrSGD in the presence of TeGS was used to establish baseline activity, while CrSGD alone served as the negative control. Reaction products were analyzed by UPLC-MS and compared to authentic standards. (A) Enzymatic assay chromatograms are shown on the same scale as an EIC at *m/z* 353. Gray signals correspond to controls in which MDRs were boiled prior to addition to TeGS assays. Geissoschizine peaks are highlighted in red and heteroyohimbanes in blue. (B) Quantitative comparison of enzymatic product formation (geissoschizine and tetrahydroalstonine). Error bars represent mean ± standard deviation. Different letters indicate statistically significant differences (pairwise Wilcoxon test, FDR corrected, p < 0.05, n = 5 biological replicates). (C) Genomic cluster of MDRs including TeGS. (D) Reaction products were reanalyzed by UHPLC-MS under optimized conditions enabling isomer separation.

### The clustered MDRs trigger GS activity and impact geissoschizine stereochemistry

While co-incubation of TeGS with specific MDRs enhanced yohimbane production, we next investigated whether these MDRs could reciprocally modulate the intrinsic activity of TeGS. We focused on the four MDRs genomically clustered with TeGS and repeated the biochemical assays under conditions enabling both qualitative and quantitative comparison of product formation (**Fig. 3A**). In the absence of TeGS, no product formation was detected. As previously observed, TeGS alone catalyzed the formation of both geissoschizine and THA in moderate amounts after 20 min of reaction. Remarkably, co-incubation with Tel01G.714, Tel01G.715, Tel01G.716, or Tel01G.717 led to a marked increase in geissoschizine and THA production, corresponding to approximately 9.3-, 2.2-, 3.8-, and 3.9-fold enhancements, respectively, relative to TeGS alone (**Fig. 3B**). A similar increase was also specifically observed for geissoschizine (**Supplemental Fig. 4A**), whereas no stimulation was detected when heat-denatured proteins were used (**Fig. 3A**). Detailed quantification further revealed a particularly pronounced effect of Tel01G.714 on THA accumulation despite the absence of detectable heteroyohimbane synthase activity for this enzyme (**Fig. 2**). This is consistent with a broad stimulatory impact on overall TeGS catalytic output. Importantly, similar trends were observed when CrGS was used in place of TeGS, indicating that the stimulatory effect of Tel01G.714, Tel01G.715, Tel01G.716, and Tel01G.717 is not restricted to the *T. elegans* enzyme but may extend to GS enzymes more generally (**Supplementary Fig. 4B; 5A-C**). Given that these MDRs do not exhibit detectable GS activity on their own, these findings suggest that Tel01G.714, Tel01G.715, Tel01G.716, Tel01G.717 and Tel01G.2847 function as auxiliary proteins that enhance the intrinsic catalytic efficiency of GS.

As GS are known to catalyze the formation of both 19*E*- and 19*Z*-geissoschizine in variable proportions ^26^, we next examined the stereochemical profile of the products generated by TeGS (**Fig. 3D**). In the absence of MDRs, TeGS predominantly produced THA, 19*E*-geissoschizine and minor amounts of 19*Z*-geissoschizine, consistent with previous observations for the *Tabernanthe iboga* orthologue TiGS ^26^. The 19*E*/19*Z* geissoschizine ratio remained essentially unchanged in the presence of Tel01G.715, confirming the minor effect of this protein on TeGS catalytic properties. In contrast, co-incubation with Tel01G.714, Tel01G.716, or Tel01G.717 resulted in a moderate increase in 19*Z*-geissoschizine levels but a marked enhancement of 19*E*-geissoschizine production. The most pronounced effect was observed with Tel01G.714, consistent with its overall impact on GS activity. In addition, this specific protein combination led to the formation of mayumbine, which likely derives from TeGS-generated intermediates. Altogether, these results indicate that Tel01G.714, Tel01G.716, and Tel01G.717 not only enhance GS catalytic output but also influence the stereochemical distribution of geissoschizine isomers, favoring the production of the 19*E* form. Such modulation is consistent with the metabolic profile of *T. elegans*, in which downstream MIAs predominantly originate from the 19*E*-geissochizine scaffold, and suggests that these MDRs contribute to shaping flux through this branch of the pathway. Due to the broad effect on geissoschizine synthesis and stereochemistry, we tentatively named Tel01G.714, Tel01G.715, Tel01G.716, and Tel01G.717, Geissoschizine Scaffolding Enzymes (GSE) 1-4.

### GSE are required for MIA biosynthesis in planta

To validate the *in vivo* role of GSEs in MIA biosynthesis, we performed functional assays *in planta* using a stepwise pathway reconstitution approach up to akuammicine. For this purpose, CrSGD and TeGS were transiently expressed in *N. benthamiana*, either alone or in combination with GSE1-4 in the presence or absence of TeGO2^13^ (**Fig. 4A**). We also included Tel01G.2847, as this enzyme was shown to produce yohimbane in the presence of TeGS, and used the inactive Tel01G.69 as a negative control.

**Fig. 4.**
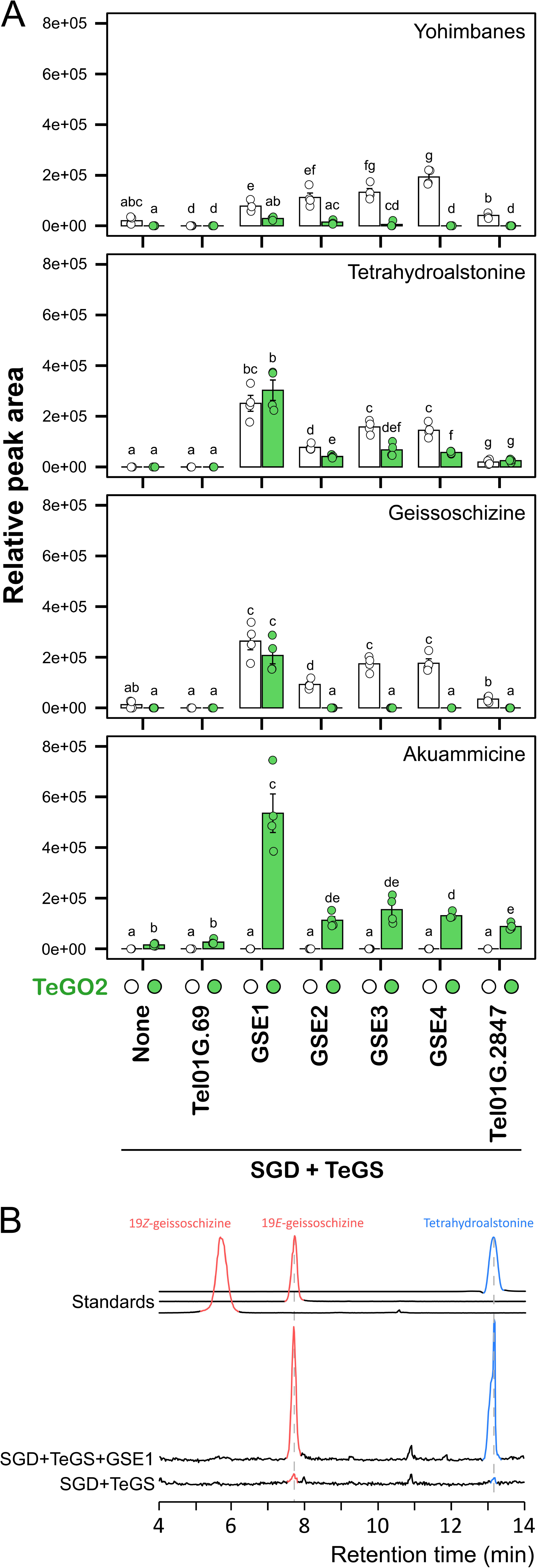
*In planta* reconstruction of the akuammicine biosynthetic pathway from strictosidine improved by GSE introduction. (A) GSE1 to GSE4 were transiently expressed in independent *N. benthamiana* leaves together with CrSGD and TeGS, in the presence or absence of TeGO2 (green and white color codes, respectively). Leaf disks were incubated with strictosidine for 24 h, and reaction products were quantified by UPLC–MS. Two additional MDRs not associated with the GS genomic cluster were also included: Tel01G.2847, which groups within the same clade as GSE2–4, and Tel01G.69, which is more divergent from the GSEs. Error bars represent mean ± standard error. Different letters indicate statistically significant differences (pairwise Wilcoxon test, FDR corrected, p < 0.05, n = 4 biological replicates). (B) Mass chromatograms (EIC at *m/z* 351) showing the separation of geissoschizine and heteroyohimbane isomers produced *in planta* in the presence of GSE1.

We first observed that the co-expression of CrSGD and TeGS led to the production of barely detectable amounts of geissoschizine, and yohimbane (PY3) but no detectable THA, slightly differing from what we observed *in vitro* (**Fig. 4A**). A similar trend was observed upon addition of TeGO2, which converted geissoschizine into only trace amounts of akuammicine (**Fig. 4A**). As expected, co-expression of GSE1-4 resulted in a dramatic increase in geissoschizine, THA, and yohimbane (PY3) biosynthesis, thus confirming the stimulatory effect previously observed *in vitro* (**Fig. 3A**; **Fig. 4A**). Interestingly, a low effect of induction was detected with Tel01G.2847 suggesting that this protein may also enhance GS activity. By contrast, no stimulatory effect was measured with Tel01G.69 indicating that the enhancing activity is restricted to a specific subset of MDRs. Overall, the most pronounced effect was again obtained with GSE1 (approximately 20.5-fold enhancement), followed by GSE3 and GSE4 that displayed comparable effects on GS activity (13.6-fold enhancement) and ended by GSE2 and Tel01G.2847. Strikingly, the geissoschizine produced under these conditions exclusively corresponded to the 19*E* isomer, suggesting that formation of the 19*Z* isomer is not favored *in planta*, as illustrated with GSE1 (**Fig. 4B**). Finally, co-expression of TeGO2 led to the conversion of geissoschizine into significant quantities of akuammicine, with production levels up to 34.5 times higher than with GS alone. This resulted in the complete consumption of the geissoschizine produced by GSE2-4 and Tel01G.2847, accompanied by a marked decrease in THA and yohimbane levels, likely reflecting a redirection of metabolic flux towards akuammicine. In contrast, although production of akuammicine was strongly increased in the presence of GSE1, geissoschizine levels remained high, suggesting that its production exceeded the conversion capacity of TeGO2. Collectively, these data confirm the strong impact of GSE1-4 as well as Tel01G.2847 on GS activity *in planta*. Consequently, Tel01G.2847 was therefore renamed GSE5.

As no gene downregulation strategy has yet been established for *T. elegans*, we searched for a potential orthologue of GSE1 in *C. roseus*, a species for which virus-induced gene silencing (VIGS) systems are well developed^27,28^. A BLAST search of the *C. roseus* genome using GSE1 identified the two close candidate genes CRO_06G024580 and CRO_06G024590. CRO_06G024580 was first amplified and cloned into the pHREAC vector for functional characterization. Biochemical assays were performed *in vitro* by combining CrSGD and CrGS in the presence or absence of the CRO_06G024580 gene product. Addition of this protein resulted in a strong enhancement of GS activity (**Supplementary Fig. 6A-B**), indicating that GSE-like proteins may also occur in other MIA-producing plants. To assess their roles *in planta* and given their high sequence identity, CRO_06G024580 and CRO_06G024590 were subsequently co-silenced in *C. roseus* cotyledons using VIGS^27^. Co-silencing of these genes led to a significant reduction in the accumulation of the two important MIAs, catharanthine and vindoline, in *C. roseus* leaves (**Supplementary Fig. 6C**). These results support a central role for GSE proteins in regulating MIA biosynthesis *in planta*.

### GSE interacts with GS and SGD

To gain insight into the molecular mechanism underlying the effect of GSEs on GS activity, we first examined the subcellular localization of GSE1–4 by transient expression in *N. benthamiana* leaves. For this purpose, each GSE and GS protein was fused to the yellow fluorescent protein (YFP) at either the N- or C-terminus, generating GSE–YFP/YFP–GSE and GS–YFP/YFP–GS constructs.

Confocal microscopy analyses were performed in transformed leaves co-expressing the different YFP fusions together with either a nucleocytosolic CFP marker or a nuclear CFP marker. These experiments revealed that GSE1–YFP/YFP–GSE1, GSE2–YFP/YFP–GSE2, and GS–YFP/YFP–GS displayed a nucleocytosolic localization (**Supplementary Fig. 7-9**). GSE3–YFP showed a comparable distribution; however, the YFP–GSE3 fusion was excluded from the nucleus (**Supplementary Fig. 10**). A similar behavior was observed for GSE4–YFP, YFP–GSE4 and GSE5– YFP, YFP–GSE5, indicating that GSE3, GSE4 and GSE5 are mainly localized in the cytosol, with a marked accumulation in the perinuclear region (**Supplementary Fig. 11-12**).

We next examined whether GS could influence the subcellular distribution of GSE proteins. When non-tagged GS was co-expressed with the GSE–YFP fusions, the localization of GSE1 and GSE2 remained unchanged. In contrast, the presence of GS resulted in a partial relocalization of GSE3, GSE4 and GSE5 to the nucleus. These observations thus suggest that interactions between GS and specific GSE proteins may occur *in planta* and prompted us to further investigate such associations (**Supplementary Fig. 13**).

To directly assess protein–protein interactions, we then performed bimolecular fluorescence complementation (BiFC) assays in *Saccharomyces cerevisiae*, complemented by co-localization analyses. BiFC experiments revealed interactions for CrSGD with GSE1-5, whereas no interaction was detected between CrSGD and TeGS (**Fig. 5A**). Consistent with these results, GSE1-4 showed nuclear or nucleocytosolic localization when expressed alone and became predominantly nuclear when co-expressed with nuclear-localized CrSGD (**Supplementary Fig. 14A, 14C, 15B**). GSE5 was an exception, showing positive BiFC signal but retaining a substantial cytosolic localization when co-expressed with CrSGD, which may reflect a weaker interaction consistent with its more limited effect on TeGS activity. In agreement with the absence of interaction in BiFC, TeGS maintained its cytosolic punctate localization, excluded from the nucleus, whether expressed alone or with CrSGD (**Supplementary Fig. 14B, 15A**). In contrast, BiFC analyses revealed clear interactions between TeGS and each GSE, which was consistent with their co-localization (**Fig. 5B, Supplementary Fig. 15C**).

**Fig. 5.**
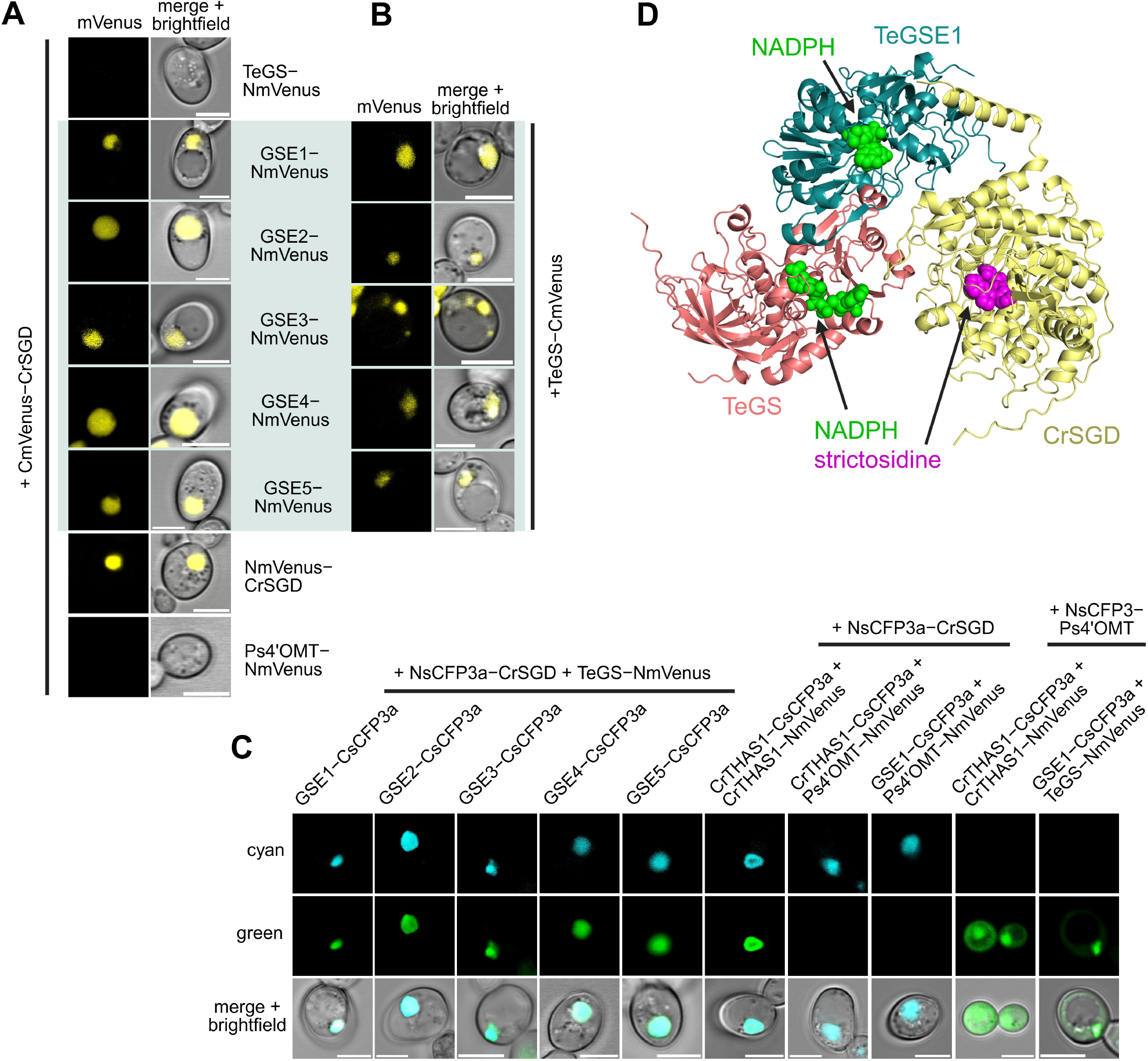
Interaction assays in yeast and ternary complex structural prediction. A) Bimolecular fluorescence complementation (BiFC) assays in *Saccharomyces cerevisiae* between C_210_mVenus-CrSGD and the indicated interaction partners. N_210_mVenus-CrSGD represents a positive control for interaction, whereas Ps4′OMT-N_210_mVenus represents a negative control. B) BiFC assays between TeGS-C_210_mVenus and the indicated interaction partners. C) Multicolor BiFC (mcBiFC) between two sets of interaction partners tagged with N_173_sCFP3a + C_155_sCFP3a (producing a cyan fluorescence signal) or N_173_mVenus + C_155_sCFP3a (producing a green fluorescence signal), previously described in ^31^. D) AlphaFold3 prediction of a ternary complex between CrSGD (yellow), TeGS (pink), and GSE1 (teal), modeled with 2 NADPH ligands (green spheres) to show the active sites of TeGS and TeGSE1. pTM=0.57, ipTM=0.45. Strictosidine (magenta spheres) was manually added to the structure to show the CrSGD active site via alignment with PDB 2JF6. Scale bars indicate 5 μm.

Given that GSE1–5 interact with both CrSGD and TeGS, we next investigated whether these interactions could occur simultaneously. To this end, we employed multicolor BiFC (mcBiFC), in which CrSGD–GSE interactions generated a cyan signal and TeGS–GSE interactions a green signal. As controls, we also included CrTHAS1, known to form homodimers and interact with CrSGD^5,6^, and an unrelated methyltransferase from *Papaver somniferum*, 3′-hydroxy-*N*-methylcoclaurine 4′-*O*-methyltransferase (Ps4′OMT), which is not expected to interact with the tested proteins ^20^. For all tested combinations, co-localized cyan and green fluorescence signals were observed, indicating the formation of ternary CrSGD–GSE–TeGS complexes (**Fig. 5C**). The specificity of these interactions was further supported by the lack of signal seen for interactions of CrTHAS1 or TeGSE1 with Ps4′OMT in mcBiFC.

To gain structural insights into the observed interactions, including a potential ternary complex, we modeled several combinations of enzyme complexes using AlphaFold3 (**Supplementary Table 3**). These included two-enzyme complexes CrSGD–GSE1 and TeGS–GSE1, as well as three-enzyme complexes CrSGD–GSE1–TeGS and CrSGD–GSE4–TeGS. We also used AlphaFold3 to predict the interactions between TeGS and GSE1 homodimers, as explained below. For each combination, we input one NADPH molecule per copy of TeGS or GSE to visualize the active sites of these MDRs. The cutoff for high-confidence predictions is usually defined as >0.5 for the overall structural confidence score (pTM) and >0.6 for the interaction confidence score (ipTM) ^29^. CrSGD– GSE1 with NADPH had scores of pTM=0.58 and ipTM=0.43. This indicates a lower confidence interface prediction; however, similar ipTM scores have been assigned to other validated transient interaction complexes^29^, so this lower score may be characteristic of low-affinity interactions. In contrast, TeGS–GSE1 with 2 NADPH had scores of pTM=0.92 and ipTM=0.92, representing a very high confidence prediction. Since many MDRs are known to form homodimers ^5,6,29^, we compared the predicted structure for TeGS–GSE1 to TeGS–TeGS (with 2 NADPH, pTM=0.79, ipTM=0.73) and GSE1–GSE1 (with 2 NADPH, pTM=0.91, ipTM=0.89) and saw extremely close alignment (**Supplementary Fig. 16B**). Indeed, TeGS–GSE1 appears to have the same interface as the homodimers, suggesting a competitive mode of binding and perhaps formation of a heterodimer between these enzymes, as suggested by BiFC experiments (**Fig. 5B, Supplementary Fig. 15C**). We further used AlphaFold3^30^ to model the three-enzyme assembly for CrSGD–GSE1–TeGS, along with 2 NADPH molecules (pTM=0.57, ipTM=0.45; **Fig. 5D**). The predicted complex positions GSE1 as a scaffold bringing the active sites of CrSGD and TeGS into close proximity, suggesting a possible role in facilitating substrate channeling between CrSGD and TeGS. In addition, a similar organization was obtained for the CrSGD–GSE4–TeGS prediction (pTM=0.57, ipTM=0.45; **Supplementary Fig. 16A**). These three-enzyme models are consistent with a hypothesis in which the GSE1–TeGS interaction forms a stable heterodimer that associates with CrSGD (**Supplementary Fig. 16C**). However, from our experimental results, we cannot rule out the possibility that GSEs might form homodimers that interact transiently with CrSGD and a TeGS homodimer, bringing them together (**Supplementary Fig. 16D**). Although the interaction confidence scores for both AlphaFold3 ternary complex models remain below the threshold for high confidence predictions, they are comparable to those reported for other validated transient interaction complexes ^29^. Therefore, the model is presented with caution as a plausible structural framework consistent with our experimental data.

Finally, co-expression of CrSGD, TeGS, and GSE proteins in yeast revealed coordinated/similar subcellular distributions of the three enzymes (**Supplementary Fig. 17**). Notably, the presence of GSEs promoted the relocalization of TeGS to the nucleus where it colocalized with CrSGD in a way that was not observed in their absence. This further supports a role for GSE proteins in mediating or stabilizing associations between pathway enzymes. Additionally, the varying degrees of co-localization between CrSGD, TeGS, and the different GSEs illustrate the highly dynamic nature of these interactions. Collectively, these results demonstrate that GSE proteins interact with both GS and SGD and support the formation of unprecedented ternary enzyme complexes, providing a mechanistic basis for their stimulatory effect on GS activity, as well as their role in coordinating MIA biosynthetic flux.

### GSEs dramatically boost MIA biosynthesis in yeast cell factories

To evaluate whether GSE proteins could represent useful tools for metabolic engineering, all GSEs, except GSE3 which is highly similar to GSE4, and Tel01G.69 were individually expressed in engineered yeast strains producing *C. roseus* or *R. tetraphylla* tryptophan decarboxylase (TDC), STR and SGD together with TeGS. All genes were cloned into CRISPR/Cas9-based expression vectors to enable stable genomic integration in yeast ^32^. The resulting transformants were grown and supplied with secologanin and tryptophan to initiate MIA biosynthesis.

In the absence of GSE expression, barely detectable amounts of geissoschizine, only reaching the detection level, were measured after 48 h of culture, consistent with previous observations highlighting the limited capacity of yeast cell factories to convert strictosidine aglycones into geissoschizine (**Fig. 6A-B**) ^33^. Under these conditions, the very low accumulation of geissoschizine may also result from partial recycling or degradation by yeast. Expression of Tel01G.69 did not affect geissoschizine production, consistent with our biochemical assays. In contrast, expression of GSE proteins resulted in enhanced geissoschizine accumulation to varying extents (**Fig. 6A-B**). GSE5 induced only a slight increase in geissoschizine levels, whereas GSE2 and GSE4 exerted a more pronounced effect. Consistent with the trends observed *in vitro* and *in planta*, GSE1 exhibited the strongest impact on GS activity, leading to an approximately 226-fold increase in geissoschizine accumulation, reaching up to 9.2 mg.L^-1^. Finally, analysis of the stereochemistry of the produced geissoschizine revealed exclusive formation of the 19*E*-isomer, as observed *in planta*, making it suitable for a downstream synthesis of valuable MIAs (**Fig. 6C**). Altogether, these results indicate that the introduction of GSE proteins, and particularly GSE1, represents a promising strategy to enhance the production of geissoschizine-derived MIAs in engineered microbial hosts.

**Fig. 6.**
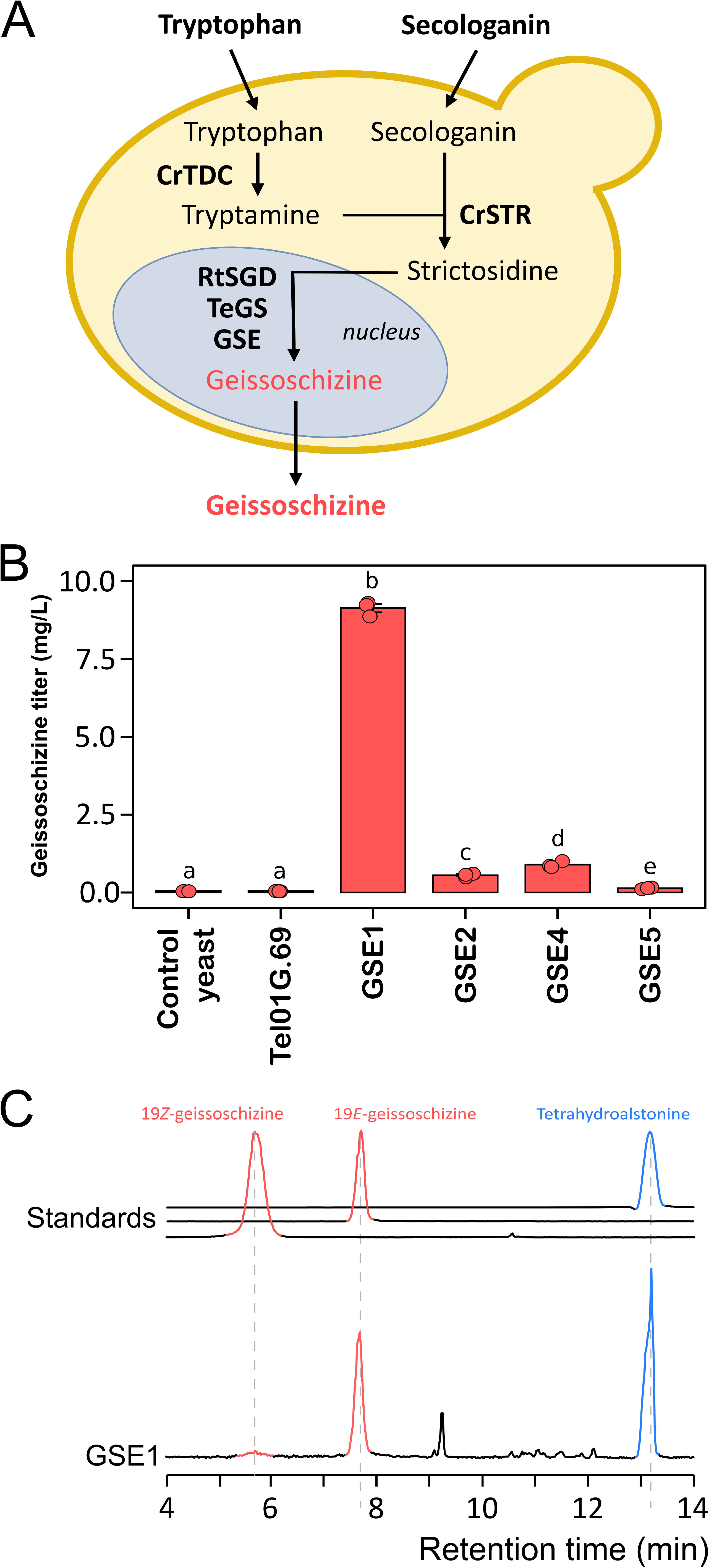
Optimization of geissoschizine bioproduction in engineered yeast using GSE. (A) Integrative yeast strains expressing CrTDC, CrSTR, RtSGD, TeGS and various GSE were fed with secologanin and tryptophan as precursors to produce geissoschizine. (B) Comparison of geissoschizine production depending on the GSE used. Control yeast did not harbor a GSE. Yeasts were grown for 48 h with precursors and geissoschizine bioconversion was quantified by UHPLC-MS. Error bars represent mean ± standard error. Different letters indicate statistically significant differences (pairwise Permutation test, FDR corrected, p < 0.05, n = 3 biological replicates). (C) Mass chromatograms showing the separation of geissoschizine and heteroyohimbane isomers produced in yeast in the presence of GSE1.

## DISCUSSION

While the biogenesis of MIAs has been investigated for several decades, the mechanisms regulating biosynthetic flux through this pathway are still not fully understood. In this context, protein–protein interactions have emerged as important regulatory determinants, as illustrated by the multimerization of SGD ^1,2^, the formation of the THAS–SGD metabolon ^5,6,20^ and enzyme assemblies involved in the final steps of catharanthine and tabersonine biosynthesis ^14,29^. Similar interaction-based regulatory mechanisms have also been described in other plant specialized metabolic pathways, including the biosynthesis of dhurrin, flavonoids, phenylpropanoid, camalexin or steroidal glycoalkaloids ^34–40^. In the present work, we reveal an additional regulatory layer by demonstrating that specific MDR proteins interact with GS, thereby affecting both catalytic efficiency and product stereochemistry.

MDR enzymes are well-known components of MIA biosynthesis, and their catalytic functions within this pathway have been extensively characterized ^3,4^.To date, however, all identified MDRs have been described as *bona fide* enzymes catalyzing defined biosynthetic steps, and no scaffolding or regulatory roles have been reported. By analyzing several *T. elegans* MDRs belonging to distinct phylogenetic clades, we uncovered an unexpected non-catalytic function for this enzyme family in controlling MIA biosynthetic flux. Through biochemical *in vitro* assays and stepwise pathway reconstitution *in planta*, we established that five MDRs (GSE1–5) exert a strong stimulatory effect on the intrinsic activity of GS, dramatically enhancing the production of both geissoschizine and THA (**Fig. 3A-D**). Importantly, none of these GSE proteins display detectable GS activity on their own, ruling out the possibility that their effect arises from redundant or cryptic catalytic functions (**Fig. 2**). While a 9.3-fold enhancement was observed *in vitro*, this effect reached up to 20.5-fold i*n planta*, suggesting a major contribution of GSEs to the overall efficiency of MIA biosynthesis (**Fig. 4A**). Among them, GSE1 displayed the strongest effect, although all GSE proteins appear to contribute to the enhancement of metabolic flux *in vivo*. Further analysis of GS reaction products revealed that, in addition to stimulating catalytic activity, GSE1–4 can also influence the stereochemistry of geissoschizine. Indeed, whereas TeGS predominantly produces 19*Z*-geissoschizine with only trace amounts of the 19*E* isomer *in vitro*, the presence of GSEs resulted in a strong shift towards the production of 19*E*-geissoschizine (**Fig. 3D**). Interestingly, this effect was even more pronounced when reactions were performed *in vivo*, as only 19*E*-geissoschizine was detected in *N. benthamiana* and yeast (**Fig. 4B; Fig. 6C**). This difference may reflect the influence of the cellular environment that could favor specific rearrangement or arise from distinct protein-protein interactions, as discussed below. Overall, this stereochemical shift is consistent with the metabolic profile of *T. elegans*, in which the major MIAs—including apparicine, dregamine, vobasine, tabernaemontaninol, and voacangine—derive from the 19*E*-geissoschizine scaffold ^41,42^.

On the other hand, GSE1–5 are not devoid of intrinsic catalytic activities. In our biochemical assays, we detected both heteroyohimbane and yohimbane synthase activities when these enzymes were combined with GS (**Fig. 2**). Notably, the heteroyohimbane synthase activity of GSE2–4 appeared restricted to the formation of THA and a putative rauniticine diastereoisomer. Interestingly, these enzymes belong to a phylogenetic clade distinct from the THAS proteins characterized in other MIA-producing plants, suggesting that they originated from an independent duplication and neofunctionalization event. By contrast, yohimbane synthesis appeared more widespread. First, we detected an intrinsic capacity of TeGS to generate a putative yohimbane compound, a property that has not been reported for other GS enzymes characterized to date ^7,26,43^. When GS was combined with GSE1-5 *in vitro*, the range of yohimbane products expanded to include several additional compounds such as rauwolscine and yohimbine, as previously reported in *R. tetraphylla*^8^. For GSE3 and GSE4, we also detected the formation of amsonine, a compound displaying an unusual C17R stereochemistry for which no biosynthetic mechanism has been previously described (**Fig. 2**) ^44^. Based on our current data, the exact origin of the yohimbane products detected in these assays remains uncertain. One possibility is that they arise from successive reduction reactions catalyzed by GS and the associated MDRs, as previously proposed ^8^. Alternatively, they may result from the enhancement of GS intrinsic activity in the presence of these proteins. Indeed, in combined assays, we observed a strong increase in the accumulation of PY3, which is typically produced solely by TeGS. Therefore, this suggests that PY3 mainly derives from enhanced GS activity.

The dual functions of GSEs as both effective auxiliary proteins and catalytic enzymes are uncommon. To date, most auxiliary proteins identified in plant specialized metabolism that can notably boost their interaction partners’ activity are non-catalytic proteins. Prominent examples include chalcone isomerase-like proteins (CHILs) that enhance naringenin chalcone production in various land plants ^45^, membrane steroid-binding proteins (MSBPs) that promote lignin synthesis in Arabidopsis^46^, and facilitator of taxane oxidation (FOTO1), which increases paclitaxel production in the Pacific yew ^47^. Meanwhile, the metabolic roles of protein-protein interactions between a scaffold enzyme and other enzymes are usually to enhance the metabolic flux towards the downstream product by recruiting the pathway enzymes into close proximity (*e*.*g*., dhurrin complex in sorghum and GAME15-centered complex in tomato^39,48^). However, these protein-protein interactions typically do not significantly increase production compared to non-catalytic auxiliary proteins. To our knowledge, GSEs are the first group of enzymes to display substantial auxiliary activity via protein-protein interaction in plant specialized metabolism.

Subcellular localization analyses combined with BiFC experiments further indicated that protein– protein interactions likely underlie the stimulatory effects of GSE proteins. While THAS and HYS have been shown to interact with SGD both *in planta* and in yeast^20^, no interaction has been reported so far between GS and SGD, also confirmed in this study (**Fig. 5A**). This absence of interaction could result in the diffusion of strictosidine aglycone intermediates toward competing biosynthetic pathways, including heteroyohimbane formation. In the present work, we demonstrate that GSE1–5 interact with GS and that a tripartite interaction between SGD, GS, and GSE proteins can also occur (**Fig. 5A-D**). Based on structural predictions of this ternary complex, we propose that GSE proteins may act as scaffolding factors connecting SGD to GS, thereby facilitating the channeling of aglycone intermediates toward geissoschizine formation (**Supplementary Fig. 16**). This could arise either from transient interactions between GSE, GS, and SGD or from the formation of a stable heterodimer between TeGS and GSEs that subsequently interacts with SGD. This latter hypothesis is supported by the differences in activity enhancement observed between *in vitro* and *in vivo* assays. Indeed, when proteins were expressed individually and next combined for *in vitro* assays, the GS activity enhancement was consistently lower (**Fig. 3A-B**). This may reflect the difficulty in disrupting pre-formed homodimers of TeGS and GSEs, thereby limiting the formation of the TeGS/GSE heterodimers. By contrast, in *N. benthamiana* leaves and in yeast, all proteins are simultaneously produced in the same cells, which likely facilitates the formation of the TeGS/GSE heterodimers and resulting in a dramatic increase in geissoschizine production (**Fig. 4A; Fig. 6A-B**). *In planta*, irrespective of the exact nature of the complexes formed, such an organization would modulate the accessibility of aglycone intermediates to competing enzymes such as THAS or HYS, thereby favoring the production of geissoschizine-derived MIAs. Crucially, this mechanism could help to explain why many MIA-producing plants contain such large quantities of geissoschizine-derived alkaloids, including vindoline and catharanthine in *C. roseus*. Such a hypothesis is also reinforced by the coexpression of TeGS and GSEs in particular with GSE1 (**Supplementary Fig. 18**).

Interestingly, the genes encoding GSE1–4 are physically clustered with TeGS in the *T. elegans* genome, further supporting a coordinated functional relationship between these enzymes (**Fig. 3C**; **Supplementary Fig. 3**). This genomic organization suggests that the potentiating function of GSE proteins may represent an additional layer of functional integration within MIA biosynthetic gene clusters. Beyond simple gene colocalization, such clusters may encode cooperative modules in which certain members evolve not solely as catalytic enzymes but also as regulatory components dedicated to the optimization of pathway flux. Notably, a similar genomic organization is conserved in *C. roseus* and other MIA-producing plants (**Supplemental Fig. 3**). Consistent with this hypothesis, GSE proteins also enhanced the activity of CrGS, and the *C. roseus* orthologue of GSE1 (CRO_06G024580) displayed a comparable stimulatory effect. Furthermore, silencing of CRO_06G024580 and CRO_06G024590 resulted in a reduction of catharanthine and vindoline accumulation in *C. roseus* cotyledons (**Supplemental Fig. 3**). Although the magnitude of this effect remained moderate—possibly due to functional redundancy among GSE proteins or to the developmental stage of the analyzed tissues—it nevertheless supports the requirement of this protein for efficient MIA biosynthesis.

Finally, expression of GSEs in yeast resulted in a strong increase in the production of geissoschizine with a notable 226-fold enhancement with GSE1 (**Fig. 6A-B**). These results further confirm the *in vivo* role of GSE proteins and open exciting perspectives for the development of efficient microbial platforms for MIA production. To date, considerable efforts have been devoted to optimizing strictosidine biosynthesis in heterologous systems, for example by improving interactions between P450 enzymes and their reductases^49^, the subsequent conversion of strictosidine aglycones into geissoschizine has consistently remained a bottleneck compared with the synthesis of heteroyohimbane and yohimbane scaffolds^11,33,50^ . The introduction of GSE proteins into engineered cell factories may therefore alleviate this limitation by promoting the efficient capture of strictosidine aglycones by GS. Ultimately, this strategy could facilitate the construction of optimized MIA biosynthetic pathways in yeast and contribute to more sustainable production routes for medically important alkaloids.

In conclusion, this work identifies a previously unrecognized regulatory role for MDR proteins in plant specialized metabolism. By enhancing both the catalytic efficiency and product stereochemistry of GS, GSE proteins emerge as key modulators of metabolic flux at a central branch point of MIA biosynthesis. Their ability to interact with different pathway enzymes, together with their conserved genomic organization across MIA-producing species, suggests that they constitute integral components of cooperative biosynthetic modules. Beyond providing new insights into the regulation and evolution of MIA pathways, our findings also highlight the potential of such auxiliary enzymes for metabolic engineering strategies aimed at improving the heterologous production of valuable plant alkaloids. From a broader perspective, this study teaches us how enzymes, traditionally considered solely as catalysts, can also evolve regulatory and flux-modulating functions, suggesting that similar metabolic boosters may remain hidden within many plant specialized metabolic pathways.

## EXPERIMENTAL PROCEDURES

### Sample sourcing

Seeds of *T. elegans* were sourced from Rarepalmseeds (https://www.rarepalmseeds.com/index.php?route=product/search&search=Toad%20tree), germinated, and transplanted into individual pots. Seedlings were cultivated at 28°C under a 16 h light / 8 h dark photoperiod. Young leaves were harvested, immediately flash-frozen, and stored at −80°C until further use.

### Microsynteny

MCscan pipeline^51^ together with the JCVI utility library (https://github.com/tanghaibao/jcvi) was used to detect and highlight microsynteny.

### Production and purification of recombinant proteins

Candidate MDR gene sequences were amplified from the genomic DNA via a nested PCR approach using Phusion High-Fidelity DNA Polymerase (ThermoScientific): initially, primers targeting the 5′ and 3′ untranslated regions of each gene were used (**Supplementary Table 4**), and the resulting amplicons served as templates for a second PCR with gene-specific primers (**Supplementary Table 4**). Purified PCR products were assembled into the pHREAC vector (Addgene plasmid #134908) or a modified pHREAC containing a BsmBI site in place of BsaI according to the method described in Besseau et al. 2025. The recombinant vectors were electroporated into *Agrobacterium tumefaciens* GV3101 cells and subsequently used for *N. benthamiana* leaf agrotransformation as described in Lezin et al. 2024. Four days post-infiltration, leaves were flash-frozen, and stored at −80°C. Frozen leaf tissues were ground in extraction buffer (50 mM NaH_2_PO_4_, 300 mM NaCl, 10 mM imidazole, 10 mM DTT, 1× EDTA-free protease inhibitor cocktail, pH 8) using a mortar and pestle. After clarification, the supernatant was incubated with TALON Metal Affinity Resin (Takara), pre-equilibrated with NPI-10 buffer (50 mM NaH_2_PO_4_, 300 mM NaCl, 10 mM imidazole, pH 8), for 2 h at 4°C under gentle agitation. Resin was washed three times with 2.5 ml NPI-20 buffer (50 mM NaH_2_PO_4_, 300 mM NaCl, 20 mM imidazole, pH 8) and proteins were eluted in four sequential steps with 500 µl NPI-250 buffer (50 mM NaH_2_PO_4_, 300 mM NaCl, 250 mM imidazole, pH 8). Eluates were desalted using Amicon 10 kDa filters (Sigma-Aldrich) with 5 ml desalting buffer (50 mM NaH_2_PO_4_, 100 mM NaCl). Protein concentrations were determined by the Bradford assay (Bio-Rad) according to the manufacturer’s instructions. Recombinant MDRs were supplemented with glycerol at 20%, flash-frozen, and stored at −80°C until use.

Separately, CrSGD was produced in *E. coli* BL21 (DE3) from pRSET-A (Invitrogen) containing the CrSGD coding sequence. The strain was grown to exponential phase until OD600 0.6, and protein expression was induced with 0.5 mM IPTG for 3 h at 30°C. Purification was achieved as described in Carqueijeiro et al., 2021.

### *In vitro* assays

For *in vitro* assays, strictosidine aglycones were generated as MDR substrates by incubating 50 μM strictosidine with 400 ng of recombinant CrSGD in 10 μL of 80 mM citrate buffer (pH 6) at 32°C and 350 rpm for 20 min. Subsequently, 1 mM NADPH, 50 mM potassium phosphate buffer (pH 7.5), and 1 μg of purified MDRs, or 1 μg of purified GS and 1µg of each MDR, were added to a final reaction volume of 100 µL. Reactions were allowed to proceed for 1 h for MDRs alone, or 20 min for GS and GSE coincubation, at 32°C and 350 rpm and were stopped by adding 100 µl of methanol. Enzymatic products were then analysed by UHPLC-MS.

### *In planta* assays

*N. benthamiana* leaf agroinfiltration was performed as described in Lezin et al. (2024)^22^. To reconstruct the pathways, different combinations of enzymes were co-expressed in the same leaves via co-infiltration of different *A. tumefaciens* strains at an OD600 of 1, mixed in equal volumes. Three days after infiltration, groups of 5 leaf disks of 1 cm diameter were collected and distributed into 24-well plates containing 750 μl of infiltration buffer with 100 μM of strictosidine. After infiltration under vacuum at 50 mBar, plates were incubated in a greenhouse for 24 h, and the medium was collected and diluted with 4 volumes of methanol prior UHPL-MS analysis. Leaves transformed with a GFP construct were used as a control.

### UHPLC-MS analyses

UHPLC-MS analyses were carried out using an ACQUITY UPLC system (Waters) coupled to a SQD2 mass spectrometer (Waters) with an electrospray ionization source. The system was controlled with MassLynx 4.2 software (Waters). Compounds were routinely separated on a Waters Acquity HSST3 C18 column (150 × 2.1 mm, 1.8 μm) at 55°C, using a flow rate of 0.4 mL/min and an injection volume of 5 μL. The mobile phase consisted of solvent A (0.1% formic acid in water) and solvent B (0.1% formic acid in acetonitrile), with a linear gradient from 10% to 50% of solvent B over 18 minutes. For improved isomer separation, compounds were injected on a Waters Acquity BEH C18 column (150 × 2.1 mm, 1.7 μm). The mobile phase consisted of solvent A (0.05% ammonium hydroxide in water) and solvent B (acetonitrile), with a linear gradient from 10% to 90% of solvent B over 18 minutes. Mass detection was performed in positive ion mode, monitoring the protonated molecular ions ([M+H]^+^).

MS/MS analyses were carried out on a Waters ACQUITY UPLC system (Waters) coupled to a Xevo TQ-S cronos mass spectrometer (Waters) equipped with an electrospray ionization (ESI) source. Chromatographic separations were performed under the same conditions as described above. Product ion spectra were obtained by targeted MS/MS fragmentation of the protonated precursor ions at m/z 353.3 (geissoschizine, heteroyohimbanes) and 355.3 (yohimbanes). The collision energy ranged from 30 V to 50 V.

### Subcellular localization studies in *N. benthamiana*

To analyze the subcellular localization of Tel01G.712, Tel01G.714, Tel01G.715, Tel01G.716 Tel01G.717, and Tel01G.2895 in *N. benthamiana*, coding sequences were amplified and cloned into pHREAC in fusion 5’ or 3’ end of the yellow fluorescent protein (YFP) coding sequence to generate YFP proteins fusions in both orientations, using primers described in **Supplementary Table 4**. The resulting vectors were electroporated into *A. tumefaciens* GV3101 as described above. Strains expressing the nucleocytosolic- and nuclear-CFP markers were also used ^52,53^. YFP fusion protein expressing strains and marker strains were prepared at OD600 of 0.05, infiltrated into *N. benthamiana* leaves. The subcellular localization was determined according to Lopez-Vazquez et al. (2026)^54^ protocol and parameters.

### BiFC and subcellular localization studies in yeast

Tel01.G714, Tel01.G716 and the codon-optimized sequence of CrSGD were synthesized by Twist Bioscience. All other *T. elegans* genes were amplified from the vectors harboring the intronless MDR sequences described above. PCR to amplify gene or backbone fragments was performed using Q5 High-Fidelity DNA Polymerase (New England Biolabs) and specific primers (**Supplementary table 4**). Enzyme-encoding genes were inserted using Gibson assembly (NEBuilder HiFi DNA Assembly Master Mix, New England Biolabs) into pre-assembled Gateway-compatible plasmids containing constitutive promoters and terminators. The gene expression cassettes were each assembled into multi-copy expression plasmids (pAG4 series) using Gateway LR Clonase II Enzyme mix (Life Technologies) (**Supplementary table 5**).

To characterize binary protein-protein interactions, the BiFC assay was developed based on the split-mVenus fluorescent protein, split at the 210th site. CrSGD was assembled into pAG425-GPD-CmV-ccdb, TeMDRs and negative control Ps4’OMT were assembled into pAG424-GPD-ccdb-NmV, and TeGS was assembled into pAG424-GPD-ccdb-NmV and pAG425-GPD-ccdb-CmV. Plasmids were transiently transformed into freshly prepared yeast competent cells (*S. cerevisiae* CEN.PK2-1D) using a Frozen-EZ Yeast Transformation II Kit (Zymo Research). Corresponding yeast strains were cultivated at 30°C 400 rpm in SD-Trp-Leu (0.17% YNB, 0.5% ammonium sulfate, 2% dextrose, and amino acid drop-out mixture) for two days. Microscopic analysis was performed using a Zeiss LSM 710 or 880 Confocal Microscope (Axio Observer, with objective Plan-Apochromat 63X/1.40 Oil DIC or Axio Examiner, with C-Apochromat 63x/1.20 W Korr) with an excitation wavelength of 514 nm and detection at 519-620 nm.

Multicolor BiFC for characterizing ternary protein-protein interactions was developed based on the design described by Waadt et al. 2008 ^31^. These constructs differed from those of our standard BiFC designs in an effort to optimize signal intensity needed for this variation of the method. In brief, the N-terminus of sCFP3a was truncated at the 173rd site, whereas the C-terminus of sCFP3a was truncated at the 155th site. Similarly, the N-terminus of mVenus was truncated at the 173rd site. CrSGD and negative control Ps4′OMT were assembled into pAG4U6-N_173_sCFP3a-GPD-ccdb, GSEs and positive control CrTHAS1 were assembled into pAG4U5-GPD-ccdb-C_155_sCFP3a, and TeGS and negative control Ps’OMT were assembled into pAG424-GPD-ccdb-N_173_mVenus. Plasmids were transiently transformed and cultured as described above. Microscopic analysis was performed as described above. Complementation of N_173_sCFP3a with C_155_sCFP3a generated cyan signal with excitation at 458 nm and detection at 462-510 nm. Complementation of N_173_mVenus with C_155_sCFP3a generated green signal with excitation at 488 nm and detection at 510-598 nm.

For co-localization studies of enzymes in yeast, CrSGD was assembled into pAG426-GPD-CFP-ccdb or pAG424-GPD-GFP-ccdb. TeGS was assembled into pAG4U6-GPD-ccdb-mCherry. GSEs were assembled into pAG424-GPD-ccdb-GFP or pAG4U5-GPD-ccdb-YFP. Plasmids were transiently transformed and cultured as described above. SYTO Deep Red Nucleic Acid Stain (Thermo Fisher Scientific) was used as a nuclear marker where indicated. Microscopic analysis was performed as described above. CFP was excited with a wavelength of 458 nm and detection at 463-556 nm (or 463-510 nm when expressed with GFP). GFP was excited with a wavelength of 488 nm and detection at 493-598 nm (or 510-598nm when expressed with CFP, or 493-512 when expressed with YFP). YFP was excited with a wavelength of 514 nm and detection at 540-588 nm when expressed with GFP and mCherry. mCherry was excited with a wavelength of 561nm and detection at 578-696 nm (or 588-696 nm when expressed with YFP).

### Structural Modeling

AlphaFold3 (alphafoldserver.com) was used to predict the structures and complexes of enzymes in combination with NADPH. Strictosidine was aligned to the active site of CrSGD using the crystal structure for SGD from *Rauvolfia serpentina* (PDB 2JF6). Predicted structures and interactions were visualized in PyMOL.

### Virus induced gene silencing in *C. roseus*

For VIGS, a 354 bp fragment of Cr06G24580 was amplified using gene-specific primers containing EcoRI and BamHI restriction sites (**Supplementary table 4**) and inserted into the multiple cloning sites of pTRV2 (Liu et al. 2002). pTRV2Cr_24580, pTRV2 and TRV1 were mobilized into *Agrobacterium tumefaciens* strain GV3101 by the freeze-thaw method. Seeds of *C. roseus* (cultivar ‘Little Bright Eye’; NESeed, USA) were used in this study. Seed germination and VIGS of target genes in *C. roseus* seedlings were performed as previously described^27^. Seedlings infiltrated with empty vectors served as controls.

For gene expression measurement, total RNA was extracted from *C. roseus* cotyledons using the RNeasy Plant Mini Kit (QIAGEN). Approximately 1 μg of total RNA was treated with DNase I to remove contaminating genomic DNA and used for first-strand cDNA synthesis with Superscript III reverse transcriptase (Invitrogen). Quantitative PCR (qPCR) was performed to measure transcript levels of the target genes with gene-specific primers (**Supplementary Table 4**). Cr*EF1α* was used as internal control for normalization.

For alkaloid extraction, *C. roseus* cotyledons were flash-frozen in liquid nitrogen, ground into a fine powder, extracted in methanol, and analyzed by UHPLC-qTOF-MS. Benzocaine was used as an internal standard to calculate the relative concentrations of the alkaloids in the samples. Authentic standards of catharanthine and vindoline (Sigma-Aldrich) were used to identify mass fragments and elution times.

### Yeast-based bioconversions

To obtain intron-less sequences of Tel01G.712, Tel01G.714, Tel01G.715, Tel01G.717, Tel01G.2895 and Tel01G.69, *N. benthamiana* leaves were agroinfiltrated with the corresponding *Agrobacterium* strains as described above. Three days post-infiltration, leaves were harvested for RNA extraction using the NucleoSpin RNA Plant and Fungi kit (Macherey-Nagel) following the manufacturer’s instructions. First-strand cDNA synthesis was performed with Maxima H Minus reverse transcriptase (ThermoScientific). All coding sequences were next amplified using Phusion High-Fidelity DNA polymerase (Fermentas) with gene-specific primers (**Supplementary Table 4**). Coding sequences of CrTDC, CrSTR and RtSGD were amplified from C. roseus cDNA. PCR products were cloned into p122, p113, p124, and p104 template vectors under constitutive promoters (**Supplementary Table 6**) using Golden Gate assembly (New England Biolabs).

Yeast strains were constructed using *Saccharomyces cerevisiae* CEN.PK113-7D (MATa MAL2-8C, SUC2) as parental strain. Expression cassettes were integrated into yeast hot-spot loci^55^ using a CRISPR–Cas9-based method ^32^. Yeast strains harbouring the pCfB2312 vector were co-transformed with gRNA helper plasmid and the appropriate NotI-linearized template vector, using the lithium acetate transformation method ^56^. Yeast transformants were selected on YPD medium plates (20 g.L^−1^ of peptone, 10 g.L^−1^ of yeast extract, and 20 g.L^−1^ of glucose) supplemented with G418 (200 µg.mL^−1^) and nourseothricin (100 µg.mL^−1^), and screened for effective integration of each expression cassette by colony PCR. Yeast strains constructed and tested in this study are listed in **Supplementary Table 6**.

For bioconversion assays, yeast were grown overnight at 30°C in 5 mL of YPD (10 g.L^−1^ yeast extract, 20 g.L^−1^ peptone, 20 g.L^−1^ glucose). The resulting cultures were used to inoculate 1 mL of fresh YPD with final concentrations of 40 µM and 200 µM of secologanine and tryptophan respectively, with an initial OD_600_ of 0,75. Cultures were incubated at 30 °C under shaking at 500 rpm. To maintain carbon and nutrient availability, cultures were supplemented with 100 µL of YP 5X and glucose 10X after 24h of growth. Samples were collected and centrifuged at 14,000 × g for 10 min to remove cells. The clarified supernatants were diluted 1:5 in methanol prior to analysis by UHPLC–MS.

### Statistical analysis

Statistical analyses were performed in R using pairwise.wilcox.test() (base package) for n > 3, and pairwisePermutationTest() (rcompanion) for geissoschizine production in yeast (n = 3). Statistical analyses are provided in **Supplementary Table 7**.

### Chemicals

Chemical used in this study include strictosidine (purified by BioCIS, Paris-Saclay, France), tetrahydroalstonine (Extrasynthese), mayumbine (Biosynth), ajmalicine (Fluka), yohimbine (Sigma-Aldrich), rauwolscine (Extrasynthese), corynanthine (Biosynth), amsonine (purified by G. Massiot, Reims, France), 19E-geissoschizine (purified by BioCIS, Paris, France), 19Z-geissoschizine (kindly provided by Pr. S. E. O’Connor, Max Planck Institute, Jena, Germany). L-tryptophan (Sigma-Aldrich) and secologanin (Phytoconsult).

## Acknowledgments

This work was supported by the Agence Nationale de la Recherche (ANR; project MIADIM – ANR-25-CE43-6607), the Région Centre-Val de Loire (ARP-IR - ScaleBio project) and the European Union’s Horizon Europe Framework (project COMBO under agreement N° 101135438). Microscope imaging data in *S. cerevisiae* was acquired through the Cornell University Biotechnology Resource Center, with NIH (S10RR025502) funding for the shared Zeiss LSM710 confocal and with NYSTEM (CO29155) and NIH (S10OD018516) funding for the shared Zeiss LSM880 confocal microscope. EPM was supported by the U.S. National Institutes of Health under award number T32GM138826.

## Author contributions

SL, CS, SB and VC conceived the project. DBB, EL, DLZ conducted the in vitro assays using recombinant enzymes. MD and CS performed all bioinformatic studies. SB and AO were responsible for GSE characterization in *Nicotiana*. SP, BP and LY performed the VIGS experiments. CBW, AL, PLP, MAB and GM performed metabolomics studies and compound identification. DBB, EL, NG, TP, NGG and NP developed yeast cell factories. EL, DBB, BSP and VC studied the GSE subcellular localization in *Nicotiana*. EPM and SL studied protein protein interaction in yeast and conducted *in silico* ternary complex modeling. DBB, EL, MD, NG, EPM, LY, SL, CS, SB and VC analyzed the data. EPM, MD, SL, CS, SB and VC wrote the manuscript with inputs of other authors.

