## Supplementary Figures for "Geissoschizine scaffolding enzymes shape monoterpene indole alkaloid biosynthesis"

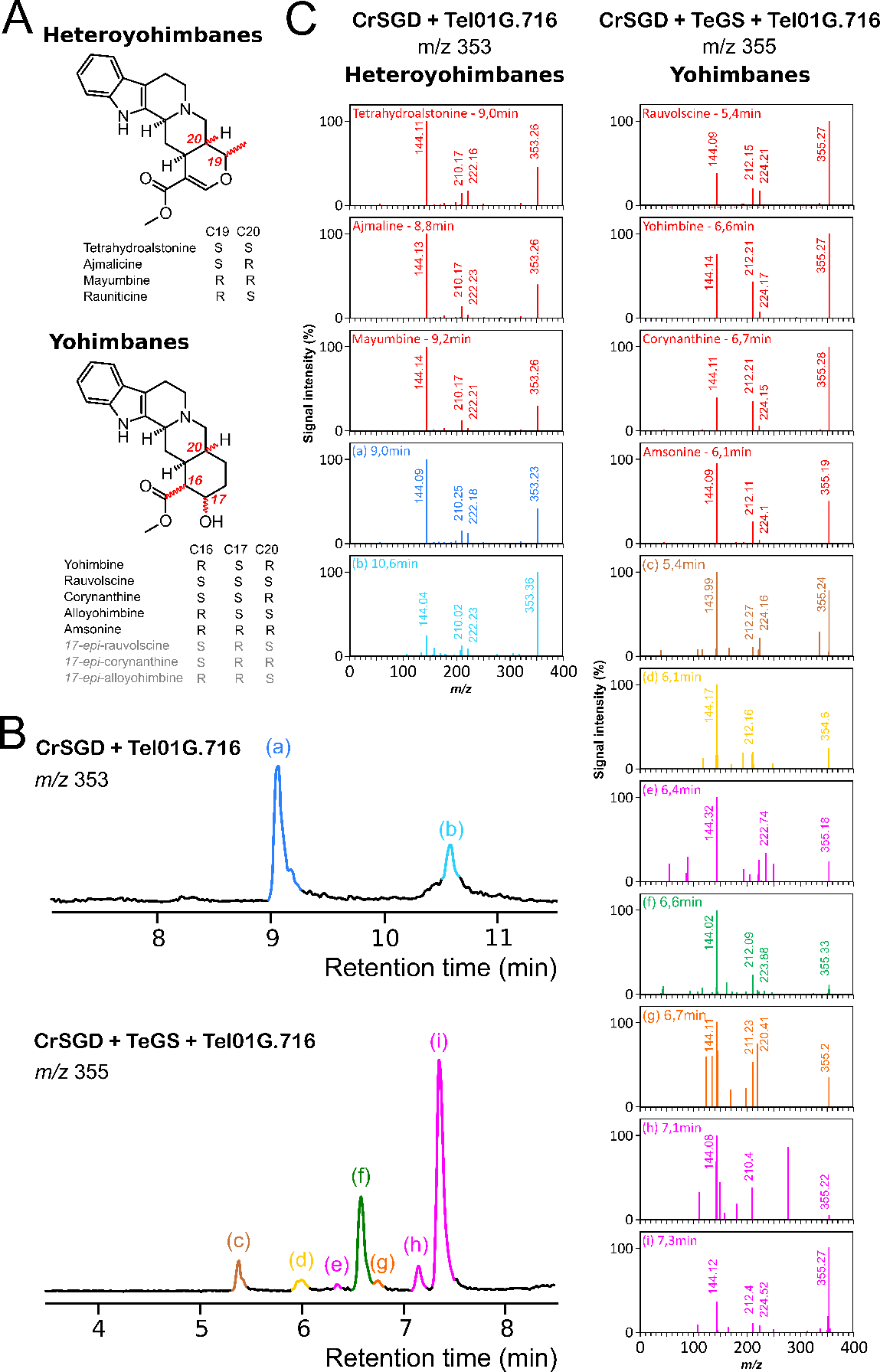

**Supplementary Fig. 1 | MS/MS fragmentation patterns of enzymatic products at m/z 353 and 355 biosynthesized by *T. elegans* MDRs.** (A) Structural diversity of heteroyohimbanes and yohimbanes (gray labels indicate predicted compounds). (B) Reaction products generated by Tel01G.716, incubated with or without TeGS, were used as references to compare fragmentation patterns with those of heteroyohimbane and yohimbane standards (red panels) shown in (C). Annotation: (a) tetrahydroalstonine, (b) putative heteroyohimbane 1, (c) rauwolscine, (d) amsonine, (e) putative yohimbane 1, (f) yohimbine, (g) corynanthine, (h) putative yohimbane 2, (i) putative yohimbane 3.

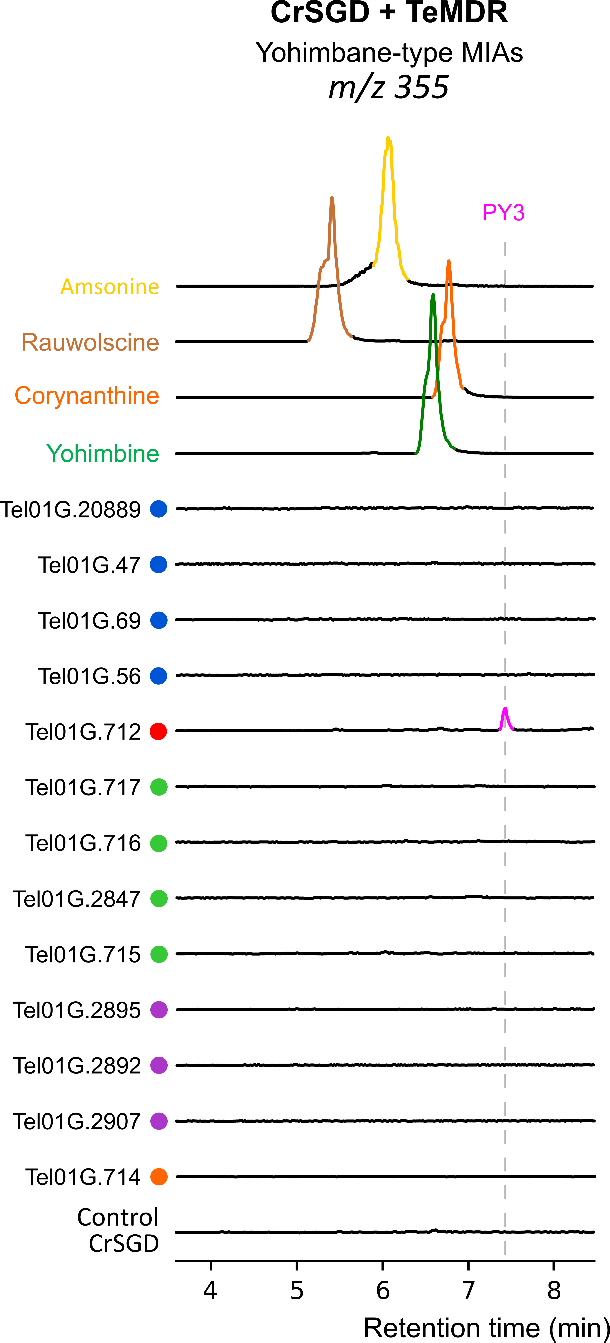

**Supplementary Fig. 2 | Screening of *T. elegans* MDR candidates by *in vitro* biochemical assays for direct biosynthesis yohimbane-type MIAs from strictosidine aglycones.** Strictosidine was pre-incubated with purified recombinant SGD from *C. roseus* followed by the addition of recombinant MDRs with NADPH. Reaction products were analyzed by UPLC-MS and compared to authentic standards at *m/z* 355. All enzymatic assay chromatograms are shown on the same scale and compared to the control condition without MDR (control CrSGD). Colored dots indicate the clade of the candidates, according to the phylogenetic tree shown in Fig. 2. PY3, putative yohimbane 3.

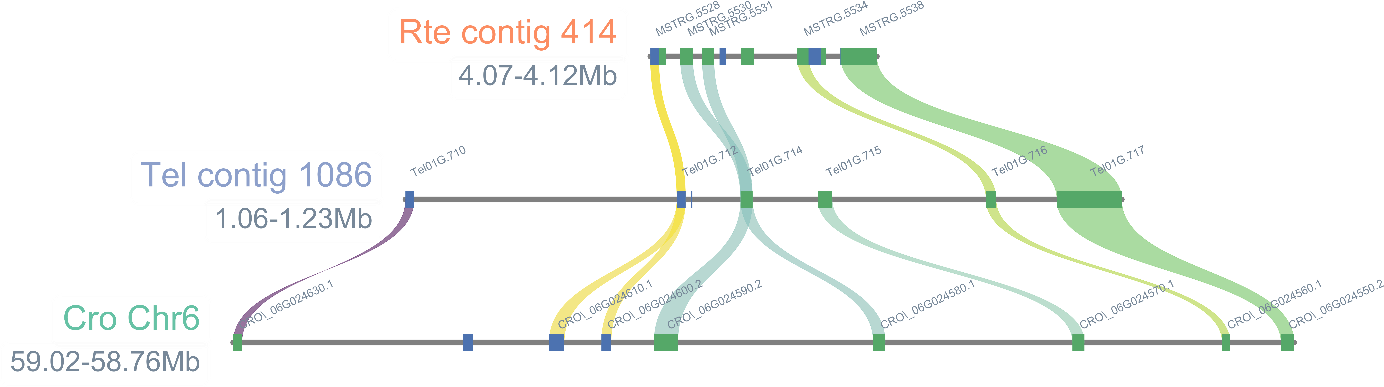

**Supplementary Fig. 3 | Micro-synteny of the *T. elegans* MDR-enriched region harboring GS across the genomes of *Rauvolfia tetraphylla* (Stander et al., 2023) and *Catharanthus roseus* (Li et al., 2023).**

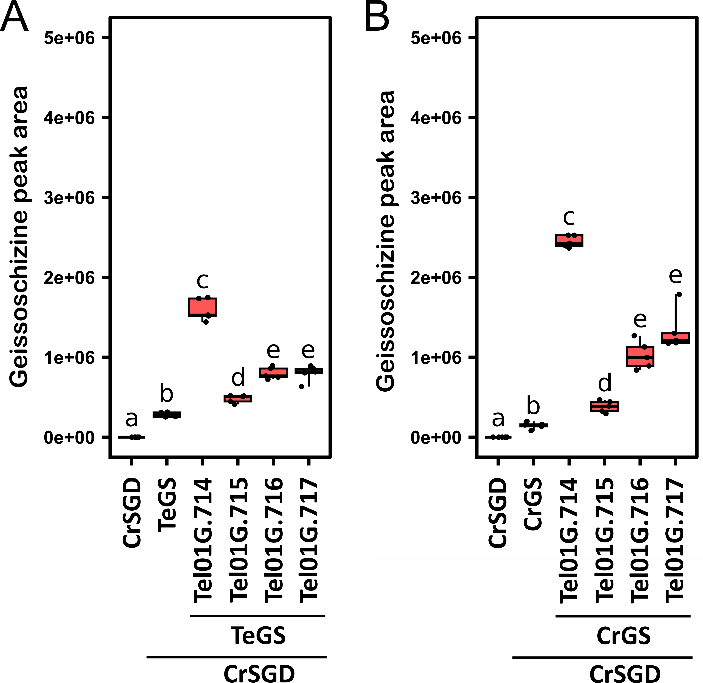

**Supplementary Fig. 4 | Quantification of geissoschizine formation in *in vitro* GS assays performed with cluster-associated MDRs.** (A) Assays using TeGS (Figure 3). (B) Assays using CrGS (Supplementary Figure 5). Error bars represent mean ± standard deviation. Different letters indicate statistically significant differences (pairwise Wilcoxon test, FDR-corrected, p < 0.05, n = 5 biological replicates).

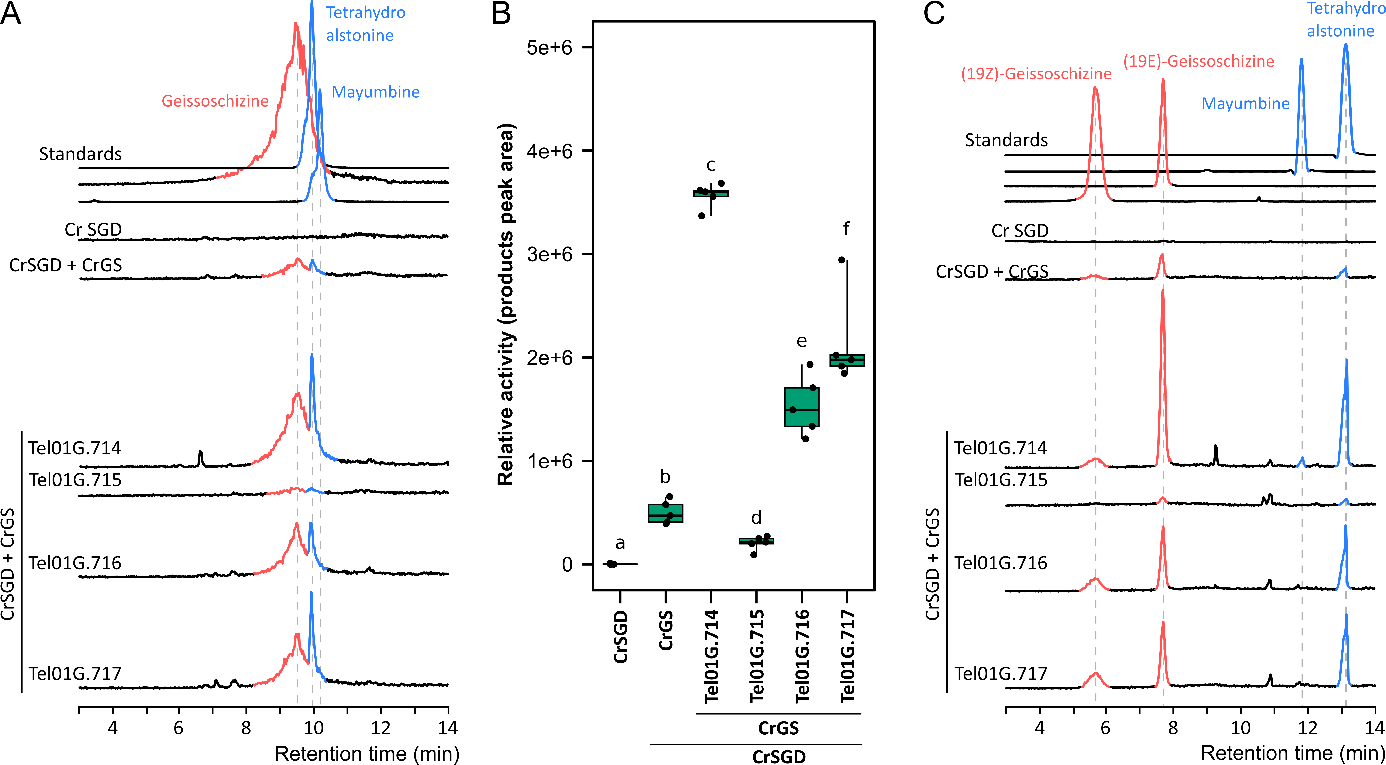

**Supplementary Fig. 5 | *In vitro* effect of cluster-associated MDRs on CrGS activity in geissoschizine and heteroyohimbane synthesis.** Strictosidine was pre-incubated with purified recombinant CrSGD, followed by the addition of CrGS together with Tel01G.714, Tel01G.715, Tel01G.716, or Tel01G.717. CrSGD in the presence of CrGS was used to establish baseline activity, while CrSGD alone served as the negative control. Reaction products were analyzed by UPLC-MS and compared to authentic standards. (A) Enzymatic assay chromatograms are shown on the same scale at m/z 353. Gray signals correspond to controls in which MDRs were boiled prior to addition to CrGS assays. Geissoschizine peaks are highlighted in red and heteroyohimbanes in blue. (B) Quantitative comparison of enzymatic product formation (geissoschizine and tetrahydroalstonine). Error bars represent mean ± standard deviation. Different letters indicate statistically significant differences (pairwise Wilcoxon test, FDR corrected, p < 0.05, n = 5 biological replicates). (C) Reaction products were reanalyzed by UPLC-MS under optimized conditions enabling isomer separation.

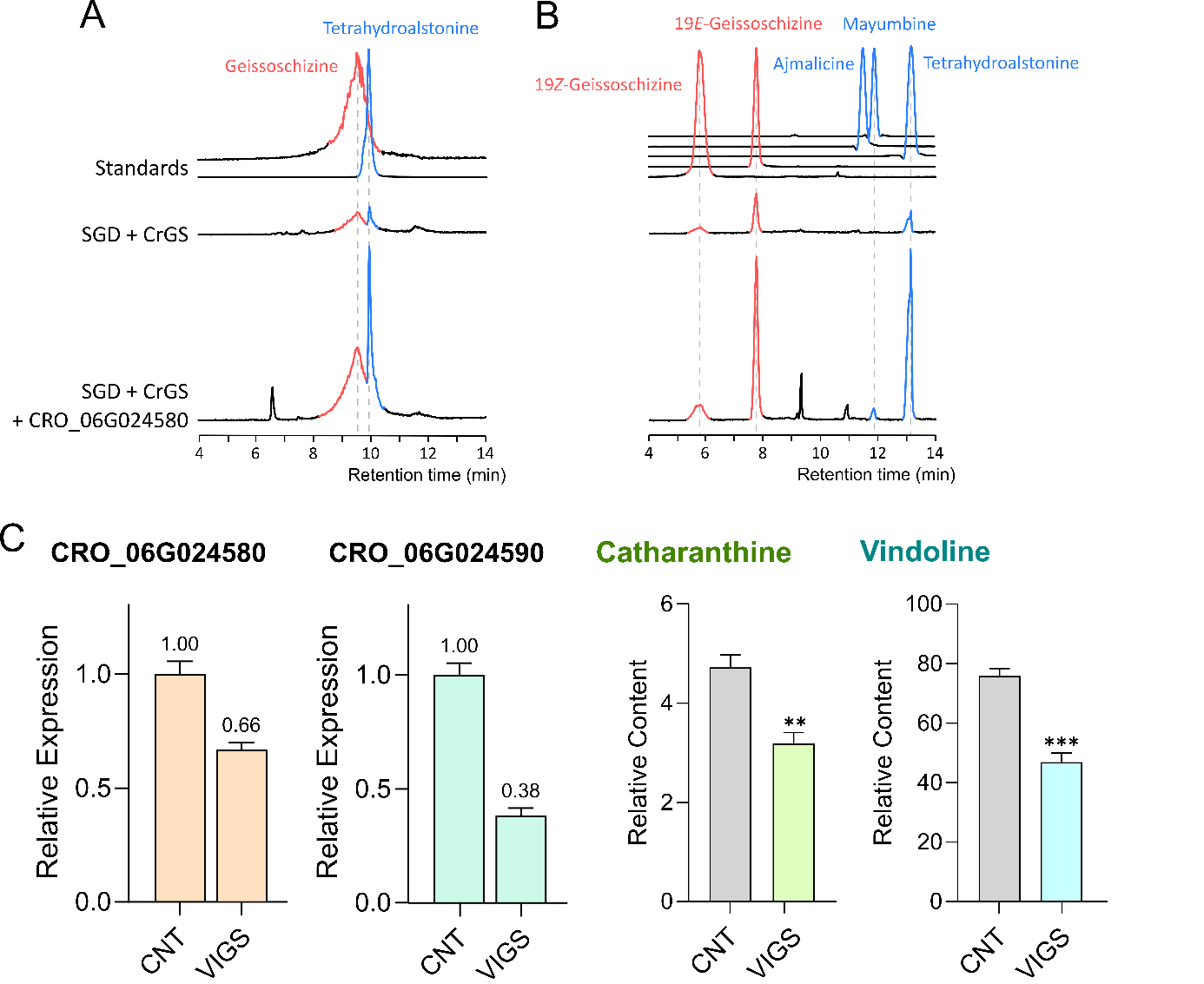

**Supplementary Fig. 6 | Functional validation of GSE *in planta* using VIGS in *C. roseus*.** (A,B) *In vitro* biochemical assays of the GSE1 orthologue from *C. roseus* (CRO_06G024580). Reaction products were analyzed by UPLC–MS using standard conditions (A) and conditions optimized for the separation of geissoschizine and heteroyohimbane isomers (B). Enzymatic assay chromatograms are shown at *m/z* 353 on the same scale. (C) *C. roseus* plants subjected to VIGS targeting CRO_06G024580/CRO_06G024590 were compared with control plants treated with the empty vector (CNT). Gene expression levels were assessed by RT–qPCR, and MIA content (catharanthine and vindoline) was quantified by UPLC–MS. Error bars represent mean ± standard error. Student’s *t*-test was used for calculating statistical significance: **, *P <* 0.01, ***, *P <* 0.001 ; n = 3 biological replicates).

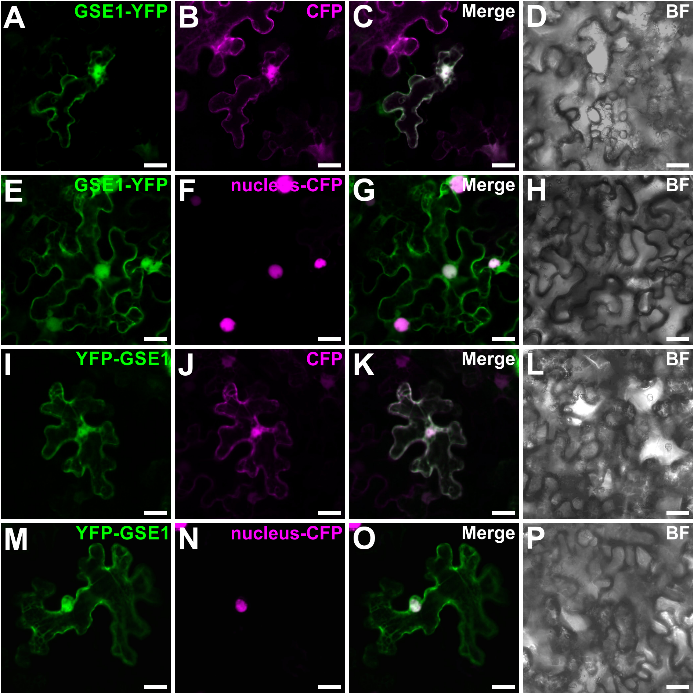

**Supplementary Fig. 7 | Subcellular localization of GSE1.** Epidermal cells of *N. benthamiana* were transiently transformed with the GSE1-YFP (**A, E**) or YFP-GSE1 construct (**I, M**) together with a nucleocytosolic CFP marker (**B**, **J**) or a nuclear CFP marker (**F**, **N**). Co-localization was confirmed after merging the two fluorescence signals (**C**, **G, K, O**). Cell morphology (**D**, **H, L, P**) was observed by bright field (BF). Bars = 10 μm.

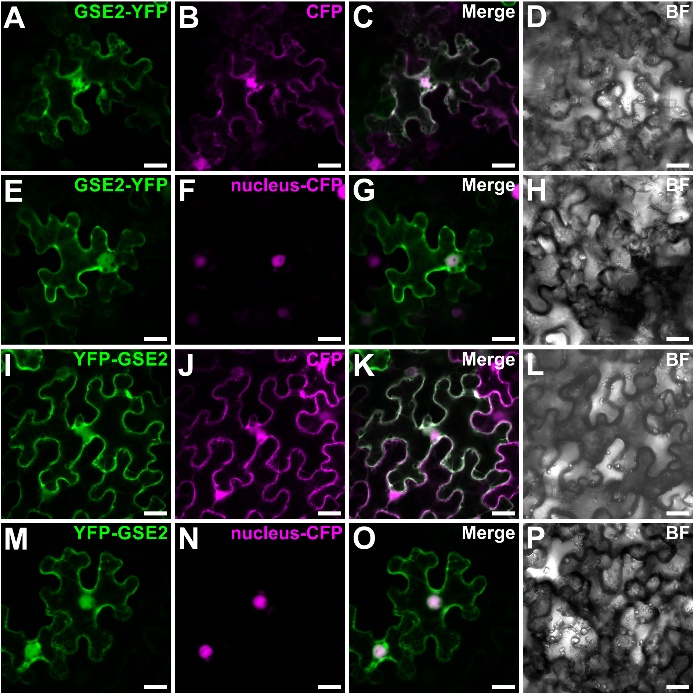

**Supplementary Fig. 8 | Subcellular localization of GSE2.** Epidermal cells of *N. benthamiana* were transiently transformed with the GSE2-YFP (**A, E**) or YFP-GSE2 construct (**I, M**) together with a nucleocytosolic CFP marker (**B**, **J**) or a nuclear CFP marker (**F**, **N**). Co-localization was confirmed after merging the two fluorescence signals (**C**, **G, K, O**). Cell morphology (**D**, **H, L, P**) was observed by bright field (BF). Bars = 10 μm.

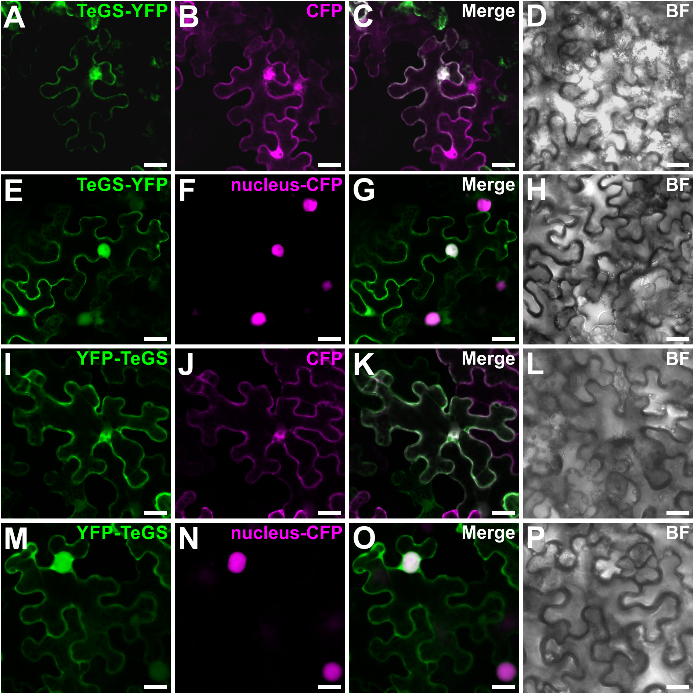

**Supplementary Fig. 9 | Subcellular localization of TeGS.** Epidermal cells of *N. benthamiana* were transiently transformed with the TeGS-YFP (**A, E**) or YFP-TeGS construct (**I, M**) together with a nucleocytosolic CFP marker (**B**, **J**) or a nuclear CFP marker (**F**, **N**). Co-localization was confirmed after merging the two fluorescence signals (**C**, **G, K, O**). Cell morphology (**D**, **H, L, P**) was observed by bright field (BF). Bars = 10 μm.

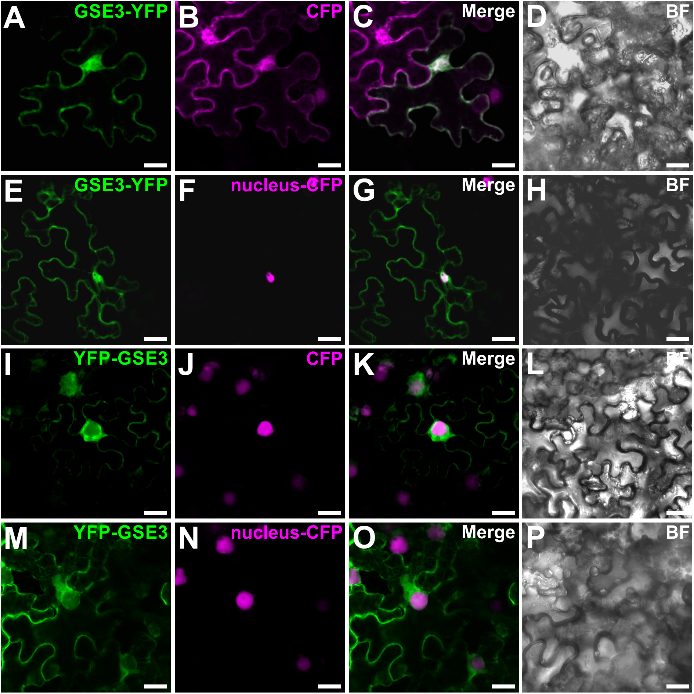

**Supplementary Fig. 10 | Subcellular localization of GSE3.** Epidermal cells of *N. benthamiana* were transiently transformed with the GSE3-YFP (**A, E**) or YFP-GSE3 construct (**I, M**) together with a nucleocytosolic CFP marker (**B**, **J**) or a nuclear CFP marker (**F**, **N**). Co-localization was confirmed after merging the two fluorescence signals (**C**, **G, K, O**). Cell morphology (**D**, **H, L, P**) was observed by bright field (BF). Bars = 10 μm.

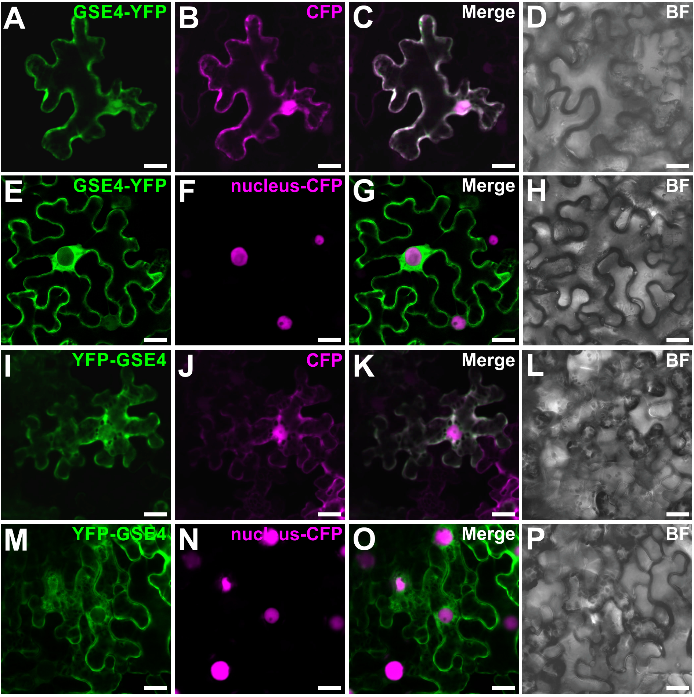

**Supplementary Fig. 11 | Subcellular localization of GSE4.** Epidermal cells of *N. benthamiana* were transiently transformed with the GSE4-YFP (**A, E**) or YFP-GSE4 construct (**I, M**) together with a nucleocytosolic CFP marker (**B**, **J**) or a nuclear CFP marker (**F**, **N**). Co-localization was confirmed after merging the two fluorescence signals (**C**, **G, K, O**). Cell morphology (**D**, **H, L, P**) was observed by bright field (BF). Bars = 10 μm.

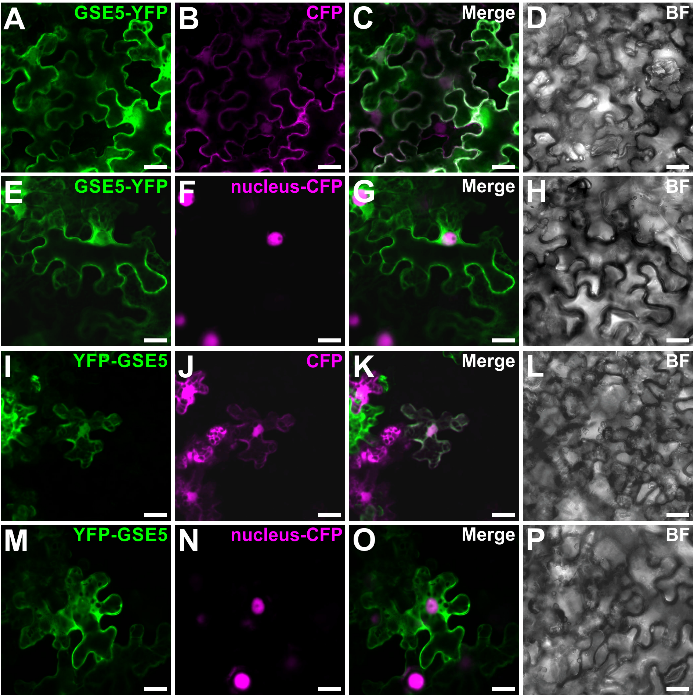

**Supplementary Fig. 12 | Subcellular localization of GSE5.** Epidermal cells of *N. benthamiana* were transiently transformed with the GSE5-YFP (**A, E**) or YFP-GSE5 construct (**I, M**) together with a nucleocytosolic CFP marker (**B**, **J**) or a nuclear CFP marker (**F**, **N**). Co-localization was confirmed after merging the two fluorescence signals (**C**, **G, K, O**). Cell morphology (**D**, **H, L, P**) was observed by bright field (BF). Bars = 10 μm.

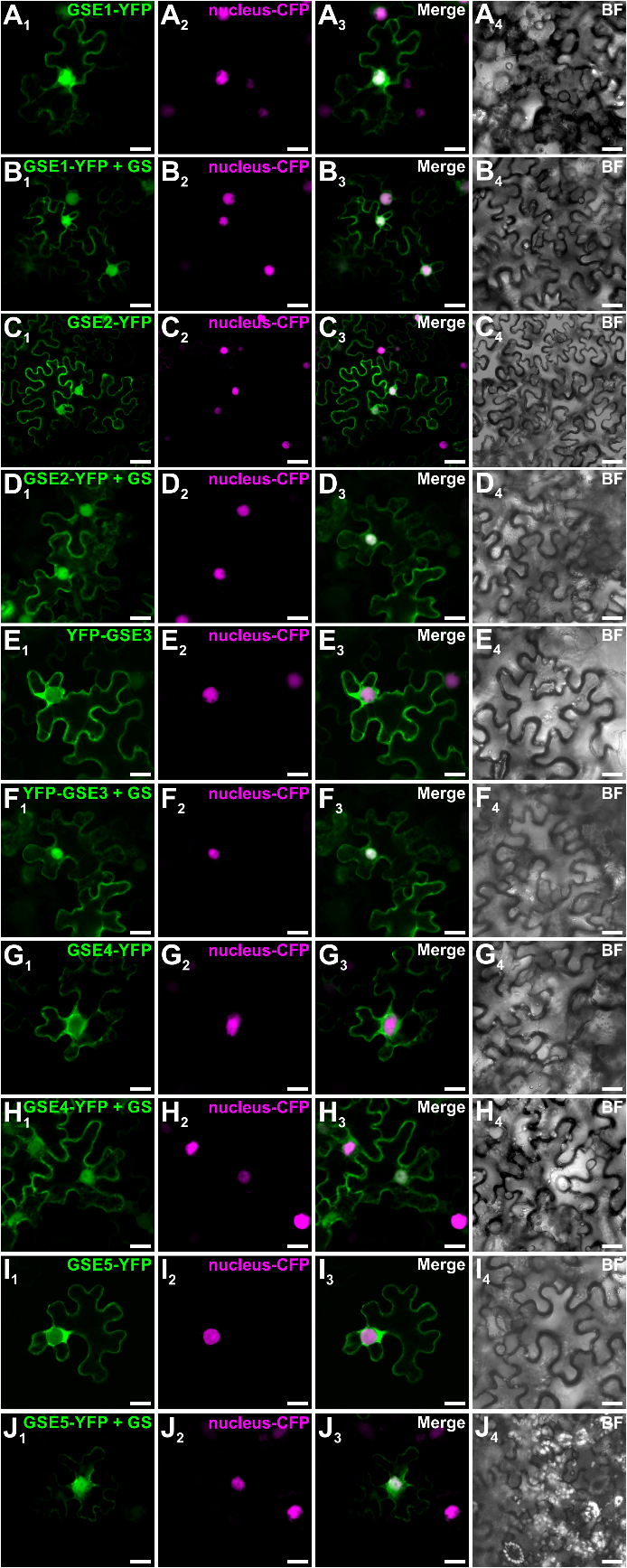

**Supplementary Fig. 13 | Impact on the coexpression of TeGS on the subcellular localisation of the different GSEs.** Epidermal cells of *N. benthamiana* were transiently transformed with the GSE1-YFP (**A1, B1**), GSE2-YFP (**C1, D1**), YFP-GSE2 (**E1, F1**), GSE4-YFP (**G1, H1**) or GSE5-YFP (**I1, J1**) alone (**A1, C1, E1, G1, I1**) or with unfused TeGS (**B1, D1, F1, H1, J1**) together with a a nuclear CFP marker (**A2-J2**). Co-localization was confirmed after merging the two fluorescence signals (**A3-J3**). Cell morphology (**A4-J4**) was observed by bright field (BF). Bars = 10 μm.

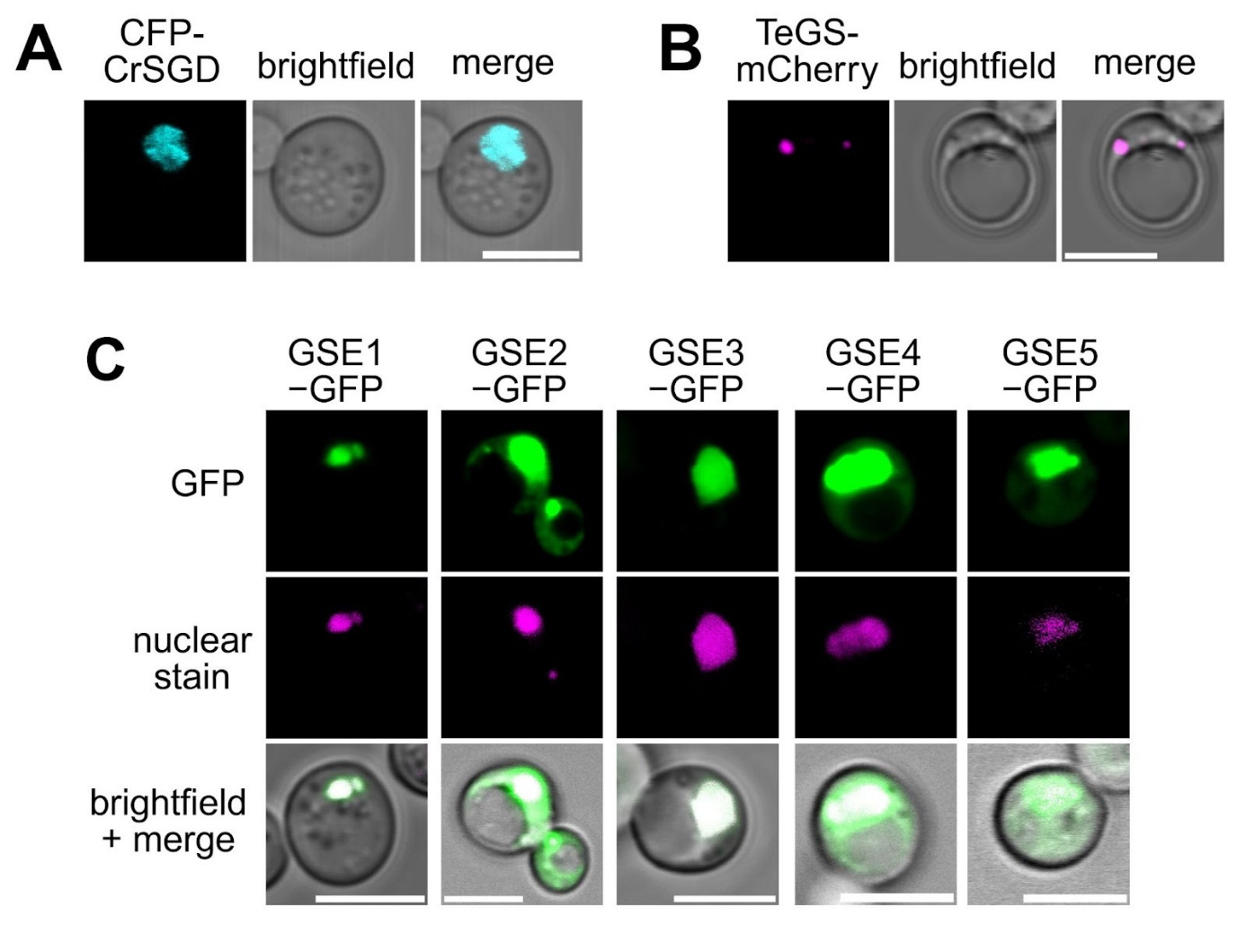

**Supplementary Fig. 14 | Localization patterns of enzymes when expressed alone in *S. cerevisiae*.** A) CFP-CrSGD, B) TeGS-mCherry, and C) GSE-GFP enzyme fusions investigated in this work. Nuclear staining used is SYTO™ Deep Red Nucleic Acid Stain. Scale bars = 5 μm.

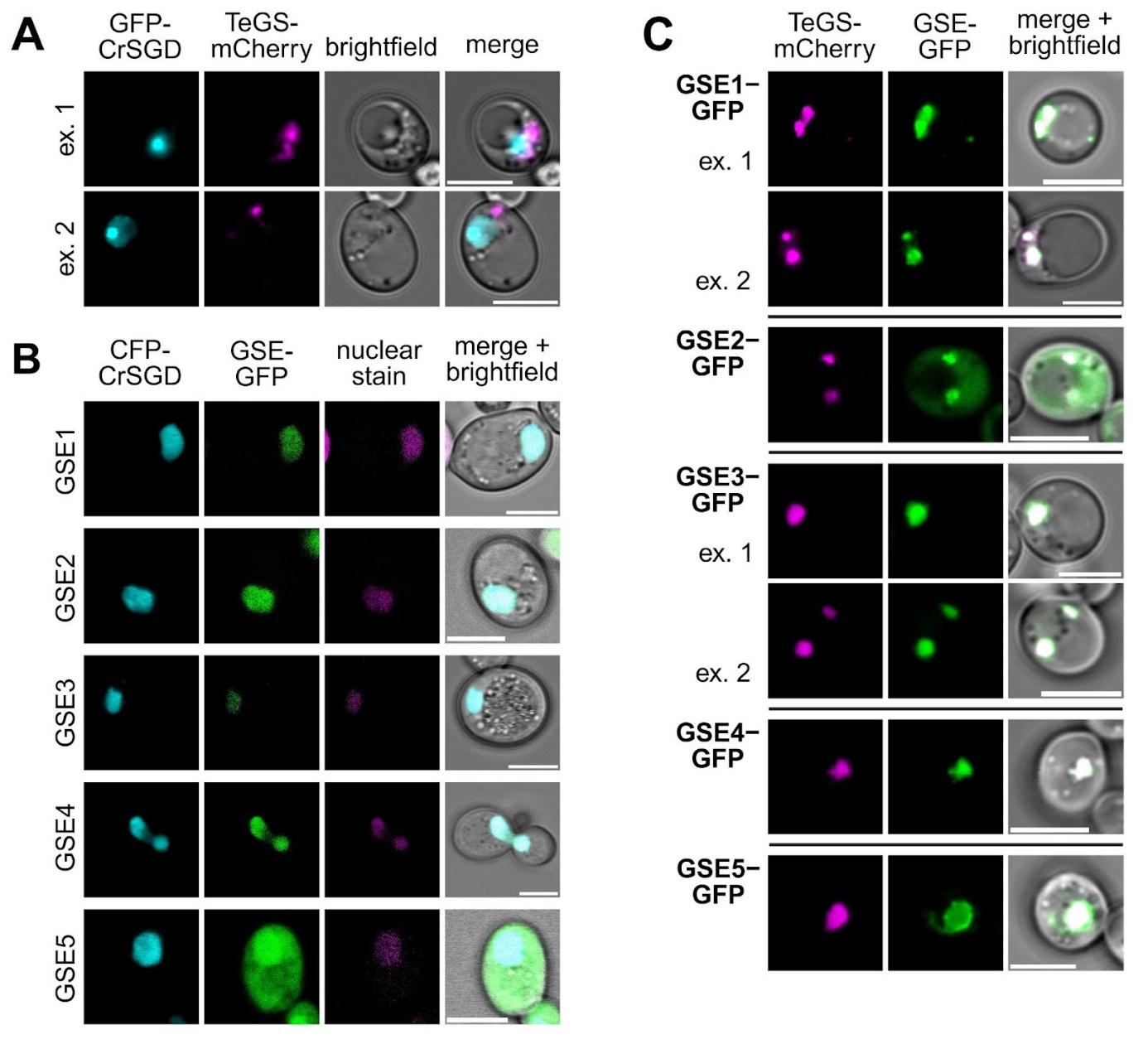

**Supplementary Fig. 15 | Localization patterns of two enzymes when co-expressed in *S. cerevisiae.*** A) GFP-CrSGD and TeGS-mCherry, B) CFP-CrSGD and GSE-GFP with SYTO™ Deep Red Nucleic Acid Stain, and C) TeGS-mCherry and GSE-GFP. Multiple examples are shown in some cases to demonstrate the variety of localization patterns observed. Scale bars = 5 μm.

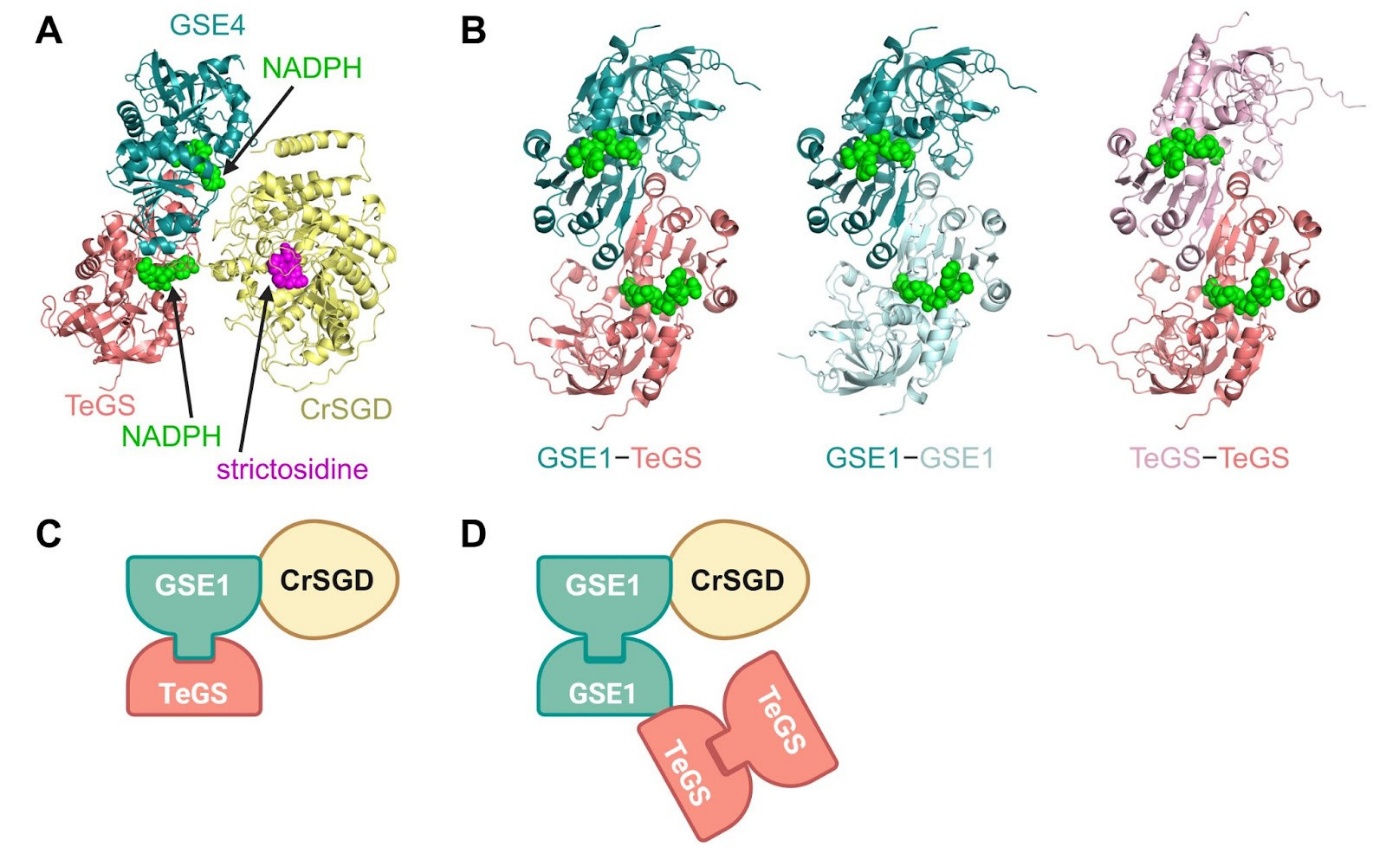

**Supplementary Fig. 16 | Comparison of *in silico* structural predictions and hypothesized multi-enzyme complexes.** A) AlphaFold3 prediction of a ternary complex between CrSGD (yellow), TeGS (pink), and GSE4 (teal), modeled with 2 NADPH ligands (green spheres) to show the active sites of TeGS and TeGSE4. pTM=0.57, ipTM=0.45. Strictosidine (magenta spheres) was manually added to the structure to show the CrSGD active site via alignment with PDB 2JF6. B) Comparison of AlphaFold3 predictions for GSE1–TeGS (pTM=0.92, ipTM=0.92, teal and pink, respectively), GSE1–GSE1 (pTM=0.91, ipTM=0.89, two-toned teal), and TeGS–TeGS (pTM=0.79, ipTM=0.73, two-toned pink). NADPH ligands are shown as green spheres. C) Schematic representation of a hypothesized three-enzyme complex in which GSE1 forms a stable heterodimer with TeGS and then transiently associates with CrSGD. D) Schematic representation of a hypothesized five-enzyme complex in which GSE1s form a stable homodimer which transiently associates with a TeGS stable homodimer and CrSGD. Made with BioRender.

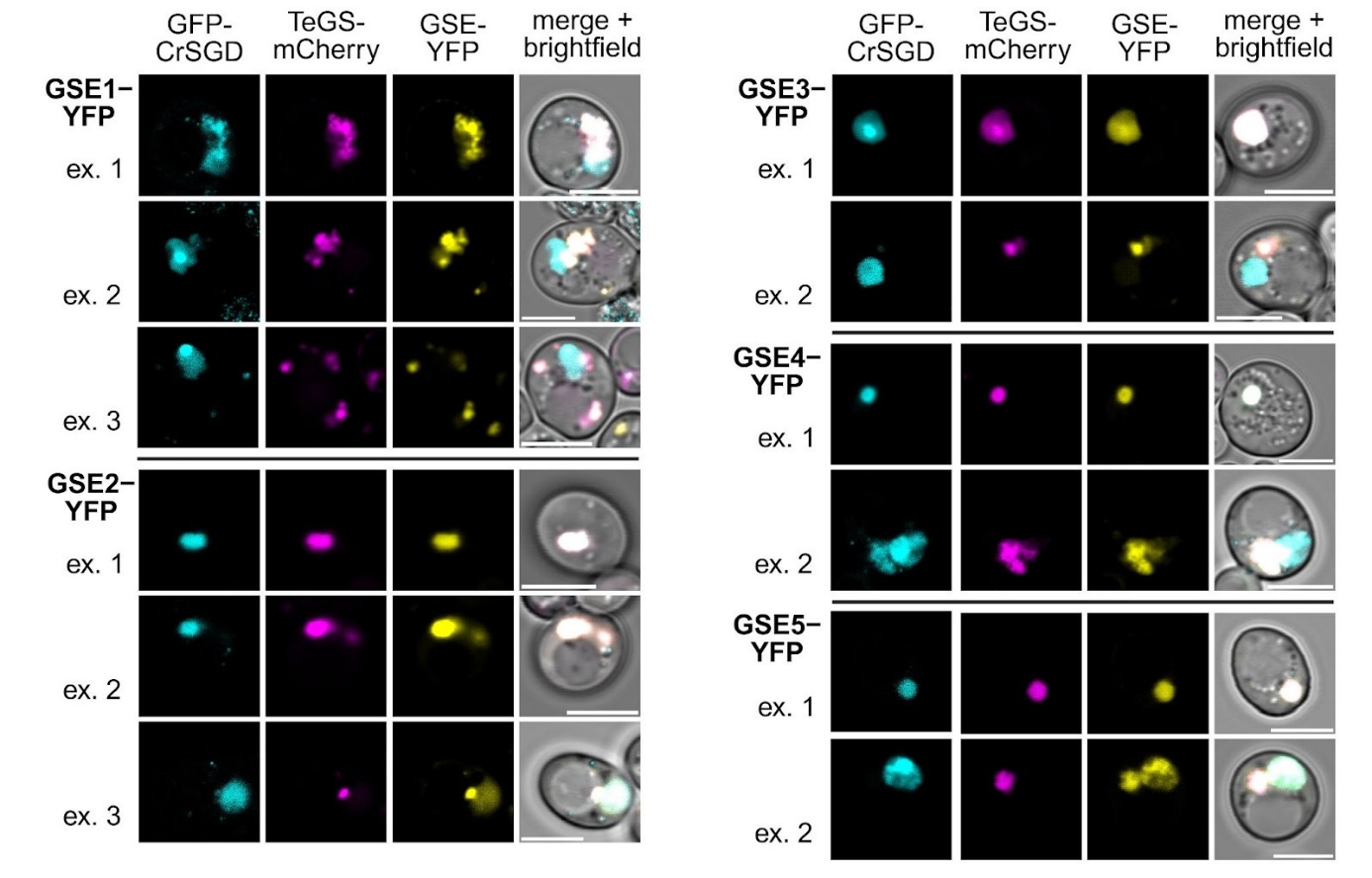

**Supplementary Fig. 17 |** **Localization patterns of 3 enzymes when co-expressed in *S. cerevisiae*.** GFP-CrSGD, TeGS-mCherry, and GSE-YFP. Multiple examples are shown to demonstrate the variety of localization patterns observed. Scale bars = 5 μm.

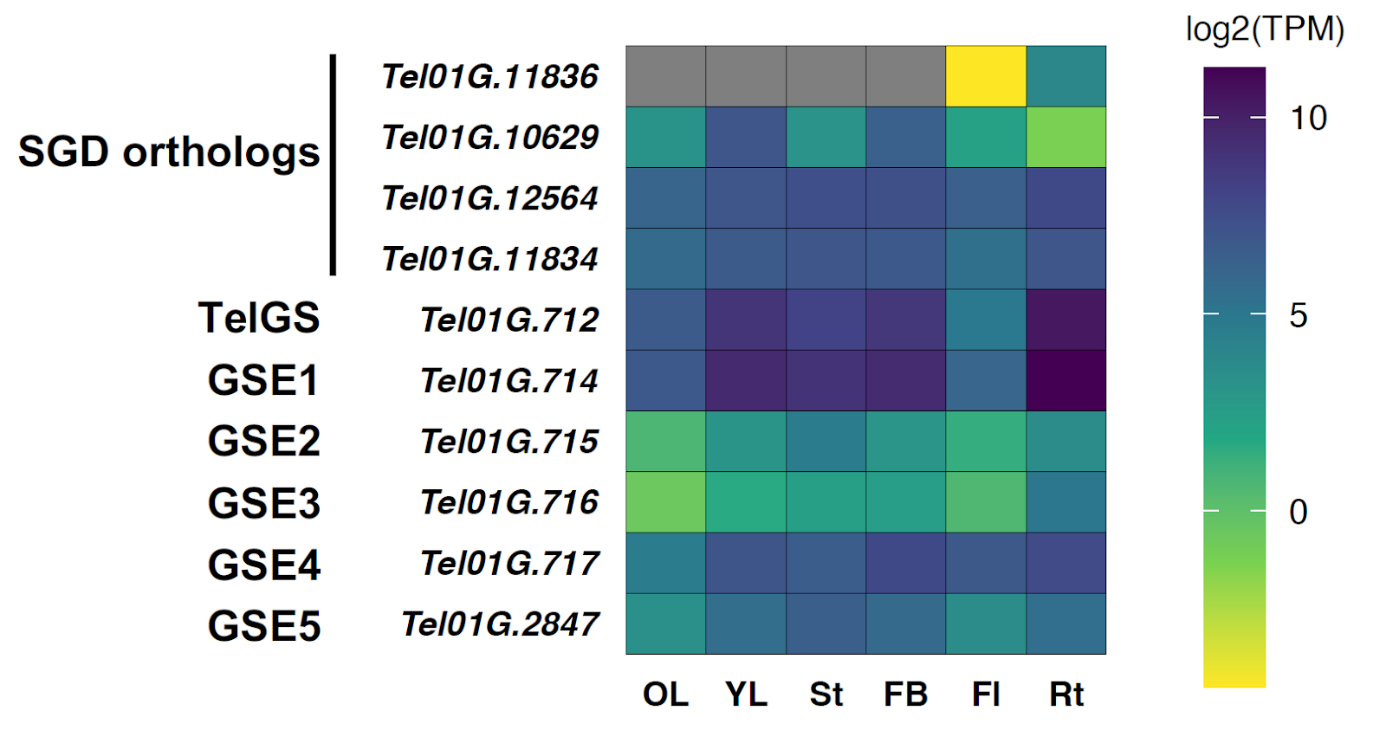

**Supplementary Fig. 18 |** **Gene expression of the predicted SGD from *T. elegans*, TeGS and GSE1-5.** OL, old leaf; YL, young leaf; St, stem; FB, floral bud; Fl, flower; Rt, root.
